# Decreased SNCA Expression in Whole-Blood RNA Analysis of Parkinson’s Disease Adjusting for Lymphocytes

**DOI:** 10.1101/2024.11.18.623684

**Authors:** Kayla Xu, Ivo Violich, Elizabeth Hutchins, Eric Alsop, Mike A. Nalls, Cornelis Blauwendraat, J. Raphael Gibbs, Mark R. Cookson, Anni Moore, Kendall Van Keuren-Jensen, David W. Craig

**Affiliations:** Integrated Translational Sciences, Beckman Research Institute, City of Hope, Duarte, CA; Department of Translational Genomics, Keck School of Medicine, University of Southern California, Los Angeles, CA; Laboratory of Neurogenetics, National Institute on Aging, National Institutes of Health, Bethesda, MD, USA; DataTecnica International, Glen Echo, MD, USA; Laboratory of Neurogenetics, National Institute on Aging, National Institutes of Health, Bethesda, MD, USA; Center for Alzheimer’s and Related dementias, National Institutes of Health, Bethesda, MD USA

## Abstract

Blood-based RNA transcriptomics offers a promising avenue for identifying biomarkers of Parkinson’s Disease (PD) progression and may provide mechanistic insights into the systemic biological processes underlying its pathogenesis beyond the well-defined neurodegenerative features. Previous studies have indicated an age-dependent increase in neutrophil-enriched gene expression, alongside a reduction in lymphocyte counts, in individuals with PD. These immune cell changes can obscure disease-relevant transcriptomic signals. In this study, we performed differential expression (DE) analysis of whole-blood RNA sequencing data from PD cohorts, incorporating a correction for immune cell-enriched gene expression, particularly neutrophil-related pathways, to improve the resolution of PD-associated molecular changes. Using 1,254 Parkinson’s Progression Markers Initiative (PPMI) samples with complete blood count (CBC) data, we developed a predictive model to estimate neutrophil percentages in a 6,987 PPMI and Parkinson’s Disease Biomarkers Program (PDBP) samples. We mitigated the confounding effects of immune cell-enriched gene expression by integrating predicted neutrophil percentages as a covariate in DE analysis. This approach revealed a consistent and significant downregulation of SNCA across all PD cohorts, a finding previously obscured by immune cell signatures. Lowered SNCA expression was found in individuals with known predisposition genes (e.g., SNCA, GBA, LRRK2) and in non-genetic PD cohorts lacking known pathogenic mutations, suggesting it may represent a key transcriptomic hallmark of the disease.

## Introduction

Parkinson’s Disease (PD) is one of the fastest-growing neurological disorders in the world, with a predicted 12 million affected individuals by 2040^1^. Compounding the global severity of PD, timely and effective treatment of the disorder remains limited by a lack of accurate biomarkers and diagnostics measures^2^. Current PD diagnosis is predominantly based on clinical features that frequently overlap with other neurological disorders, leading to high rates of misdiagnosis^2,3^. As such, reliable diagnostic biomarkers could sustainably improve patient prognosis via early detection and potential therapy targets^2,3^.

Recent advancements in PD diagnostic biomarkers include the cerebral spinal fluid (CSF) based alpha-synuclein seed amplification assay (SAA), demonstrating high sensitivity^4-6^. Blood-based transcriptomic biomarkers in PD, however, are still highly researched^7-9^. The advantages of blood-based biomarkers include the less invasive and more universally applicable nature of blood draws and tests compared to the more invasive lumbar puncture required to obtain CSF samples^7^. Analysis of transcriptomic variation in PD may also provide a better understanding of the disorder’s underlying biological mechanisms. Prior GWAS analyses have identified 90 independent risk signals explaining 16–36% of the heritable risk of PD, suggesting a significant genetic component of the disease that may be present in the transcriptome^10^. Previous studies have identified significant gene-level alterations in PD blood samples, such as in pathways related to immune activity, inflammation, mitochondrial function, cell death, etc.^9^. These studies, however, are limited by small sample sizes, which restricts statistical power and makes reproducibility of differentially expressed genes identified in each study challenging^9^.

The Parkinson’s Progressive Markers Initiative (PPMI) and Parkinson’s Disease Biomarkers Program (PDBP) are two multi-center, longitudinal observational studies developed to identify PD biomarkers^11,12^. These two datasets include whole blood RNA-sequencing data for healthy control and PD participants across multiple visits, including PD participants with known PD risk variants (i.e. SNCA, LRRK2, and GBA)^11-13^. Previously, Craig et al. evaluated RNA expression in the PPMI cohorts and found a strong enrichment of immune-related genes and pathways, specifically an upregulation of neutrophil degranulation in pathway analysis^13^. This discovery is consistent with prior work establishing a relationship between PD and immune activity, and several studies have found neutrophil count or neutrophil-to-lymphocyte ratio to be a potential PD biomarker^14^.

How neutrophils function in PD pathogenesis is still unclear. Some studies have argued PD may be caused by dysregulated inflammatory responses that trigger a-syn (SNCA) aggregates or general overexpression of a-syn in dopaminergic neurons, and that increased expression of a-syn may, in turn, increase inflammation, resulting in a cycle of a-syn aggregation that leads to neurodegeneration^15-17^. In the brain, a-syn accumulation has been linked to pro-inflammatory factors, changes in astrocyte activity, and microglia hyperactivity^18^. However, evidence is limited linking a-syn and neutrophil expression, specifically in inflammatory responses. According to the Human Protein Atlas, some blood cell types do express SNCA, including neutrophils, monocytes, and dendritic cells, but SNCA expression in neutrophils is relatively low at 9.8 pTPM compared to the highest expression of SNCA in plasmacytoid dendritic cells at 115.5 pTPM^19^.

In this study, we further evaluate the impact of neutrophil expression in PD using 3,700 PPMI and 2,790 PDBP longitudinal whole blood RNA-seq samples from 1406 PPMI and 1164 PDBP participants. Samples were obtained from healthy controls (n=1,026), individuals with idiopathic PD (n=1,054), individuals with a mutation in SNCA (n=24), LRRK2 (n=419), and GBA (n=347). Only a subset of PPMI samples (n=1,254) had complete blood counts (CBC), so a regression learning model was built to predict neutrophil percentage in the remaining samples (n=5,236). Differential gene expression analysis correcting for predicted neutrophil percentage was conducted between control and PD cohorts, and results were further analyzed to identify potentially significant pathways and gene interactions. SNCA improves as a statistically significant DE signal with neutrophil percentage as a design covariate, suggesting a transcriptomic-level suppression of SNCA in whole blood that occurs independently from neutrophil-related inflammation in PD. We further establish a potential mitochondrial gene expression signature in PD cohorts distinct from healthy control samples.

## Results

Out of 8,461 total samples, bulk short-read RNA-sequencing data of 6,987 whole blood samples from 2,711 participants in PPMI and PDBP passed filtering for analysis. Filtering criteria included QC quality metrics as well as the removal of BioFIND samples due to small sample size (see Methods for more details). 1,254 of the PPMI samples had known neutrophil percentages from corresponding complete blood count (CBC) data, and as such, were used to develop machine learning models to predict neutrophil percentage in the 5,643 PPMI and PDBP samples without CBC data. 407 of the 6,987 passing samples had a diagnosis other than ‘Case’ or ‘Control’ upon enrollment into PPMI or PDBP and were removed from downstream analysis.

### Neutrophil percentage linear modeling and prediction

To predict neutrophil percentage from whole blood gene expression counts, we developed multiple machine learning models and compared their performance to select the best model for prediction in the 5,643 samples with no known neutrophil percentage. To avoid data leakage between the test and training sets, we insured samples from the same participants were not present in the train and test set simultaneously by performing an 0.8-0.2 train test split on participant IDs, then assigning the participant samples to the corresponding set. Training set and testing set gene counts were then normalized and transformed using DESeq2 variant stabilization transformation separately, with design = ∼1 to ensure normalization was unbiased by sample metadata.

The first model we developed was a linear model based on genes known to be enriched in blood cells (Fig. 1a). Blood cell-enriched genes were selected based on the Human Blood Atlas for neutrophils, lymphocytes (T-cells and B-cells), monocytes, eosinophils, basophils, and dendritic cells^20^. From the 2,070 enriched genes present in the normalized counts, a linear model was first created for each cell type to find the genes most predictive of neutrophil percentage by cell type. Backward elimination was applied until all genes in each model had a p-value of less than 0.05. The 118 significant genes were then combined to create a final linear model, once again applying backward elimination until all remaining genes had a p-value less than 0.05. The final blood cell-based linear model contains 27 significant genes (Supplemental Table 1).

**Fig. 1.**
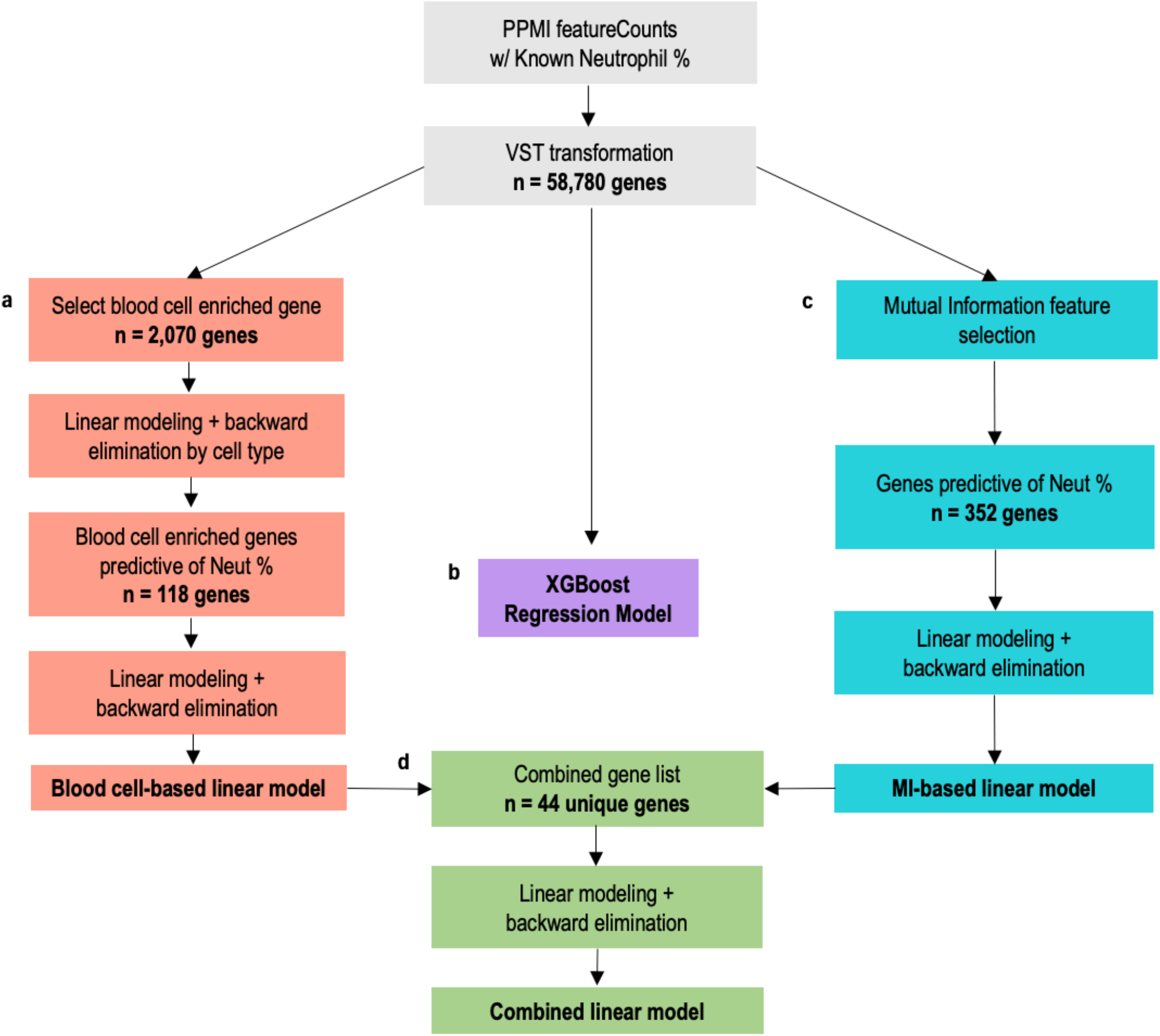
Workflow diagram of regression model development for predicting neutrophil percentage from gene expression data. 1,254 passing samples with CBC test results were used to create machine learning regression models to predict neutrophil percentage. **a**,**c**,**d**, Train-test splits for regression model development were created by randomly splitting the 600 unique participants between an 80% train set and 20% test set, then assigning the respective samples to the corresponding set. Three different linear models were created to compare the performance of different methods of feature selection: (**a**) biology-based via selection of only blood cell enriched genes, (**c**) data-driven via mutual information feature selection from all genes, and (**d**) combining the methods to include genes from both biology-based and data-driven selection. **b**, Additionally, an XGBoost regression model (**b**) was developed with all 58,780 transformed gene counts. We used the best performing model to predict neutrophil percentage for 2,932 AMP samples and 2,711 PPMI samples with no known neutrophil percentage.

The second model was a linear model based on data-driven feature selection (Fig. 1c). We used mutual information (MI) feature selection to identify which of the 58,780 total genes had the highest dependency with neutrophil percentage. We applied a MI score threshold of 0.3, resulting in 352 genes with high dependency. Like the first model, these 352 genes were then used to create a linear model, and backward elimination was applied until all p-values were less than 0.05. The final MI-based linear model contained 17 genes, of which only 2 were present in the blood cell-based model. However, all genes in the model were enriched in neutrophils, based on the Human Blood Atlas (Supplemental Table 2).

We then developed a third linear model based on the final genes in the previous blood cell-based and MI-based models (Fig. 1d). The combined 44 unique genes were used to train the combined model, once again applying backward elimination until all p-values were less than 0.05. This combined model contained 31 genes: 21 from the blood cell-based model, 8 from the MI-based model, and the 3 genes which were present in both (Supplemental Table 3)

In addition to linear modeling, we implemented a XGBoost regression model which considered all 58,780 genes to predict neutrophil percentage (Fig. 1b). XGBoost is a method of gradient tree boosting that has demonstrated improved predictive performance in many fields and applications, including expression-related prediction^21,22^. In our implementation, we used the XGBoost R package to create a regression model with hyperparameters nrounds = 10, eta = 0.3 and max depth = 3.

We evaluated and compared the four models by calculating the average R-squared, root mean squared error (RMSE), and mean absolute error (MAE) in the test set across 100 train-test splits. The worst model across all three metrics was the MI-based model, with statistically significant poor performance across all metrics. When evaluating based on the R-squared in test sets, the combined linear model significantly outperforms all other models (Fig. 2a). For RMSE, while the combined model can still be considered a top performer, there was a nonsignificant difference between RMSE values from the combined model and the cell-based model (Fig. 2b). The same is true for MAE (Fig. 2c).

**Fig. 2.**
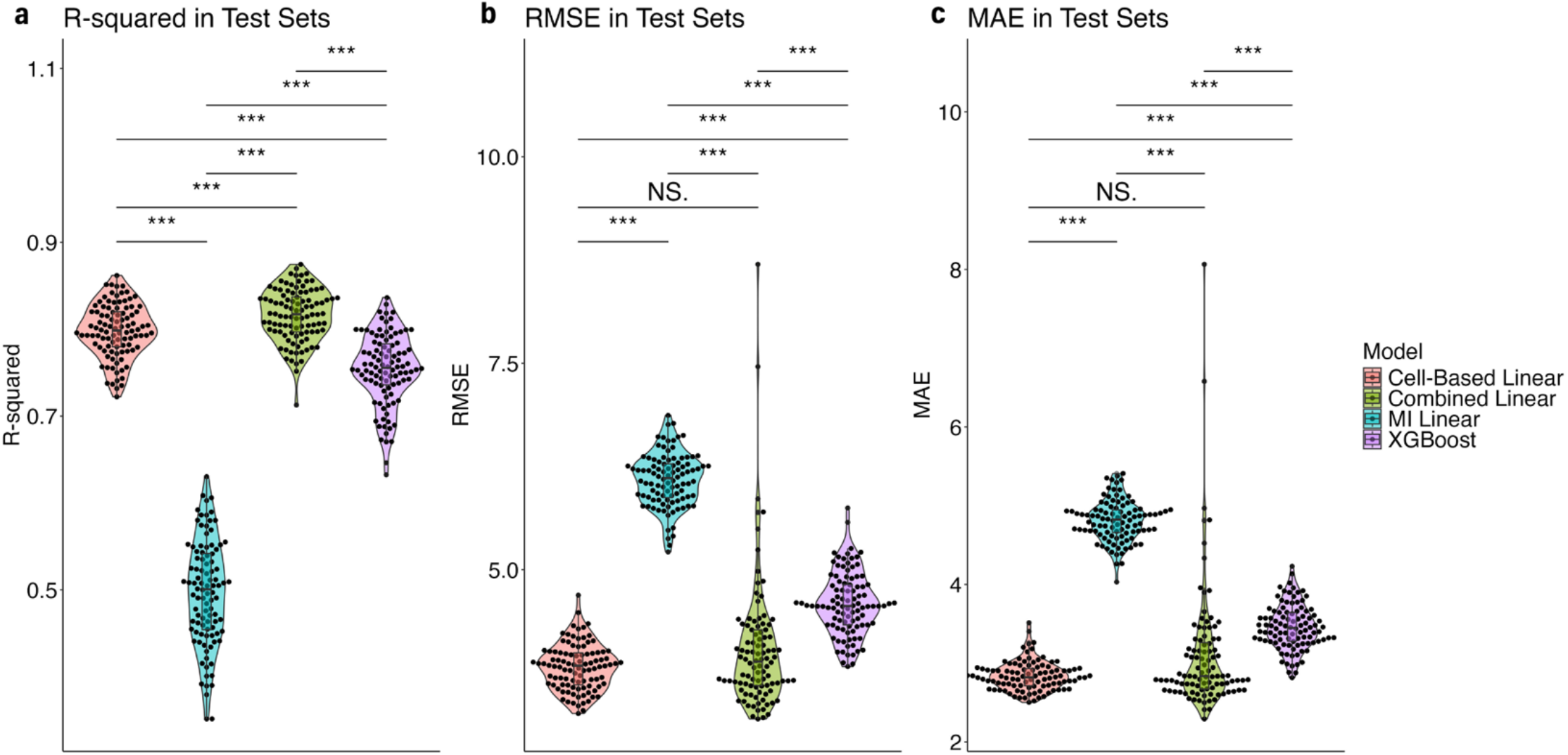
Comparison of different machine learning models to predict neutrophil percentage in PPMI and PDBP patients. **a**,**b**,**c**, Each model type was trained and tested on 100 train-test splits where samples were split 0.8-0.2 by participants. (**a**) plots the R^2^ value of each model when applied to the test sets, (**b**) plots the root mean squared errors, and (**c**) the mean absolute errors. A Wilcoxon signed-rank test was used to test the statistical significance of differences between the models for each metric. *, **, *** indicate p-values less than 0.05, 0.01. and 0.001, while N.S. indicates no significance.

As such, we selected the combined linear model as the best performer for neutrophil percentage prediction. A final linear model was fitted using all 1,254 samples and the 31 combined model genes. We then used the model to predict neutrophil percentage in the samples without CBC data. The combined list of 5,643 predicted and 1,254 known neutrophil percentages were used in all downstream analyses.

### Analysis of sample variation

Before performing differential expression analysis, we evaluated the validity of our covariate design, as well as the effect of including predicted neutrophil percentage, using PCA. PCAs were created using plotPCA() in DESeq2. We first conducted PCA on vst counts from all 6,987 samples, in which we found a very high and statistically significant correlation between PC1 and the sex of the participant (Supplemental Fig. 1a). We also saw a certain degree of correlation with sample QC metrics such percent intronic bases, percent mRNA bases, percent usable bases, etc. in PC2 to PC5. A slight correlation with percent chimeric reads was also present, and as such we included a small filter for >3% chimeric reads in the passing samples (see Methods). Predicted neutrophil percentage was most highly correlated with PC8, indicating that the neutrophil percentages predicted by our model do appear to correlate with some variation in gene expression. In a second PCA of the VST counts, this time using the limma removeBatchE5ect() function in R to correct for our selected design covariates (ie. disease status, sex, percent mRNA bases, participant age, and neutrophil percentage), we can successfully eliminate the effect of neutrophil percentage and other covariates/confounders (Supplemental Fig. 1b). Removing such effects also substantially decreased the percent variance explained by each PCA, demonstrating the positive impact of controlling for these covariates in the differential expression design matrix (Supplemental Fig. 1c,d).

### Differential gene expression analysis with predicted neutrophil percentage

We conducted differential gene expression analysis on multiple PD cohort vs control comparisons and considered the difference between analysis with predicted neutrophil percentage as a covariate and without. The full design = ∼case + sex + age squared + percent mRNA bases + predicted neutrophil percentage was determined through variance analysis and design testing (Supplemental Table 4). For each cohort comparison, the DE analyses were conducted with PPMI and PDBP samples combined, as well as with samples separated by study and at baseline (i.e. age at initial blood draw upon enrollment into PPMI or PDBP) (Supplemental Fig. 3-5).

By accounting for predicted neutrophil percentage, we see a substantial decrease in the number of differentially expressed genes in PD case vs all control samples from both PPMI and PDBP studies (Fig. 3a,b). A subset of these genes are neutrophil-enriched genes that are eliminated with correction, which indicates that inclusion of predicted neutrophil percentage as a covariate is an effective proxy in DE analysis for true neutrophil percentage (Fig. 3e,f). Neutrophil correction additionally appears to correct for all leukocyte-enriched genes, not just those in neutrophils (Supplemental Fig. 6).

**Fig. 3.**
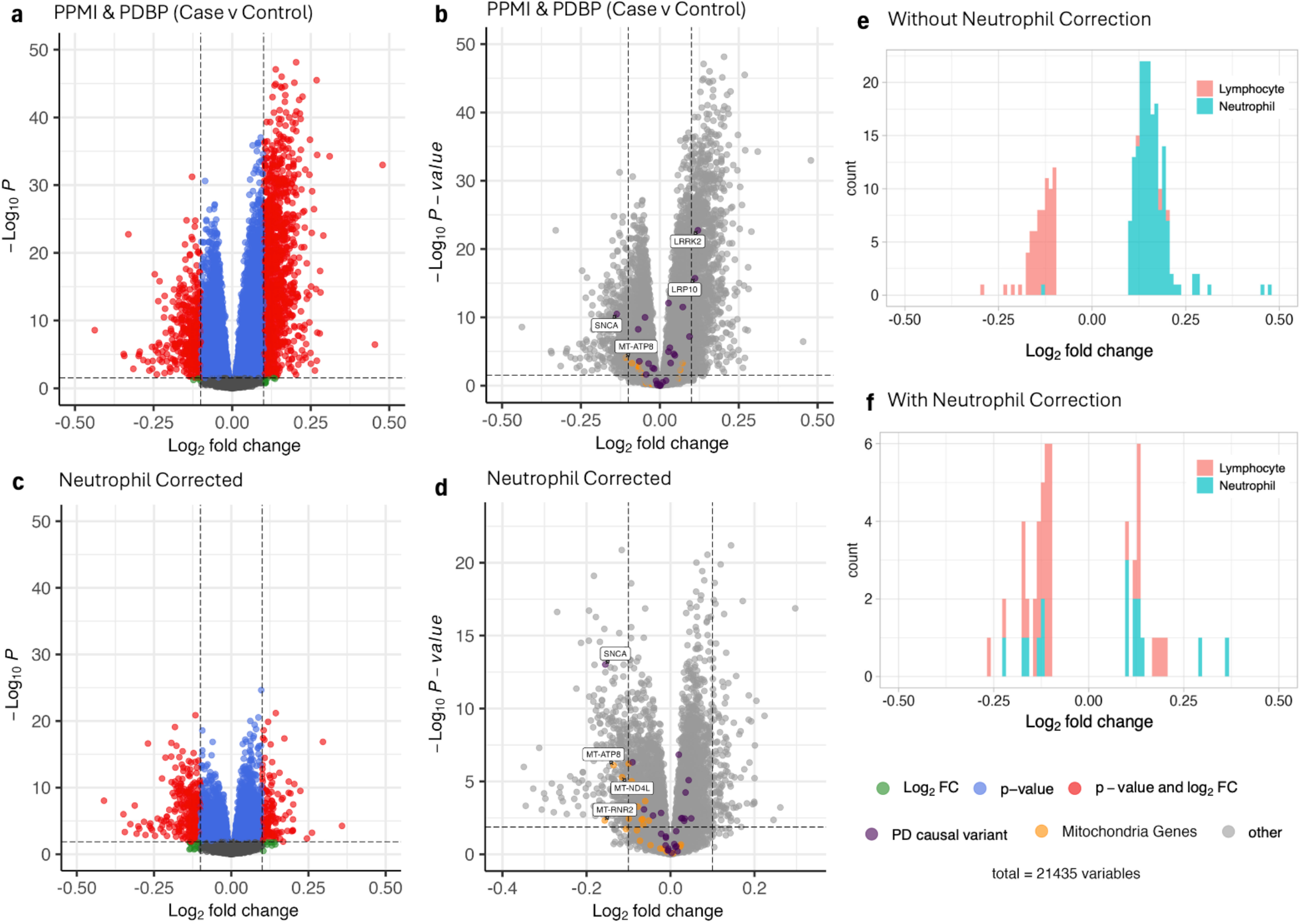
Differential expression in patients comparing case to controls, with and without controlling for predicted neutrophil percentage. **a**,**b**,**c**,**d**, Volcano plots of differential expressed genes, without controlling for predicted neutrophil percentage (**a**,**b**) and with controlling for predicted neutrophil percentage (**c**,**d**) in the design matrix. Genes with a log_2_ fold change of > 0.1 or < -0.1 are considered differentially expressed and colored in red. Genes which are either known PD causal variants or mitochondrial genes are colored purple and orange respectively in the volcano plot without controlling for predicted neutrophil percentage (**b**) and with controlling for predicted neutrophil percentage (**d**). PD causal variants and mitochondrial genes that are differentially expressed are additionally labeled by their gene name. **e**,**f**, Histograms show the distribution of significantly differentially expressed neutrophil- and lymphocyte-enriched genes in the differential expression analysis without (**e**) and with (**f**) controlling for predicted neutrophil percentage.

SNCA was the only known PD causal variant to remain differentially expressed after neutrophil correction. Not only does SNCA maintain statistical significance, but the removal of genes related to neutrophil percentage additionally improves the DE signal of SNCA: SNCA is the 973^rd^ most significant DE gene without neutrophil correction (Fig. 3b) and the 18^th^ after neutrophil correction (Fig. 3d, Supplemental Table 4). Two PD causal variants, LRRK2 and LRP10, are no longer differential expressed after neutrophil correction (Fig. 3b,d). LRRK2 was found to play a role in neutrophil chemotaxis in Mazaki et al., which may explain why neutrophil correction resulted in an extreme decrease in LRRK2 significance^23^. LRP10 has been shown to cluster with genes related to neutrophil degranulation in RNA analysis conducted by the Human Blood Atlas^20^. Overall, we can see that neutrophil correction eliminates the DE signals of blood cell-enriched or related genes, while simultaneously improving the DE signal of SNCA in our PD case vs control comparison. This statistically significant depression of SNCA is also present when plotting log(CPM) of the gene counts, corrected for predicted neutrophil percentage (Fig. 4f, Supplemental Figure 14). We additionally establish that SNCA expression doesn’t appear to correlate with our predicted neutrophil percentage, suggesting that the mechanisms behind lower SNCA expression in PD whole blood occurs independently from immune cell-related activity (Supplemental Fig. 15). The lack of correlation is further corroborated by the increased significance of SNCA downregulation in DE analyses after neutrophil correction (Fig. 3).

**Fig. 4.**
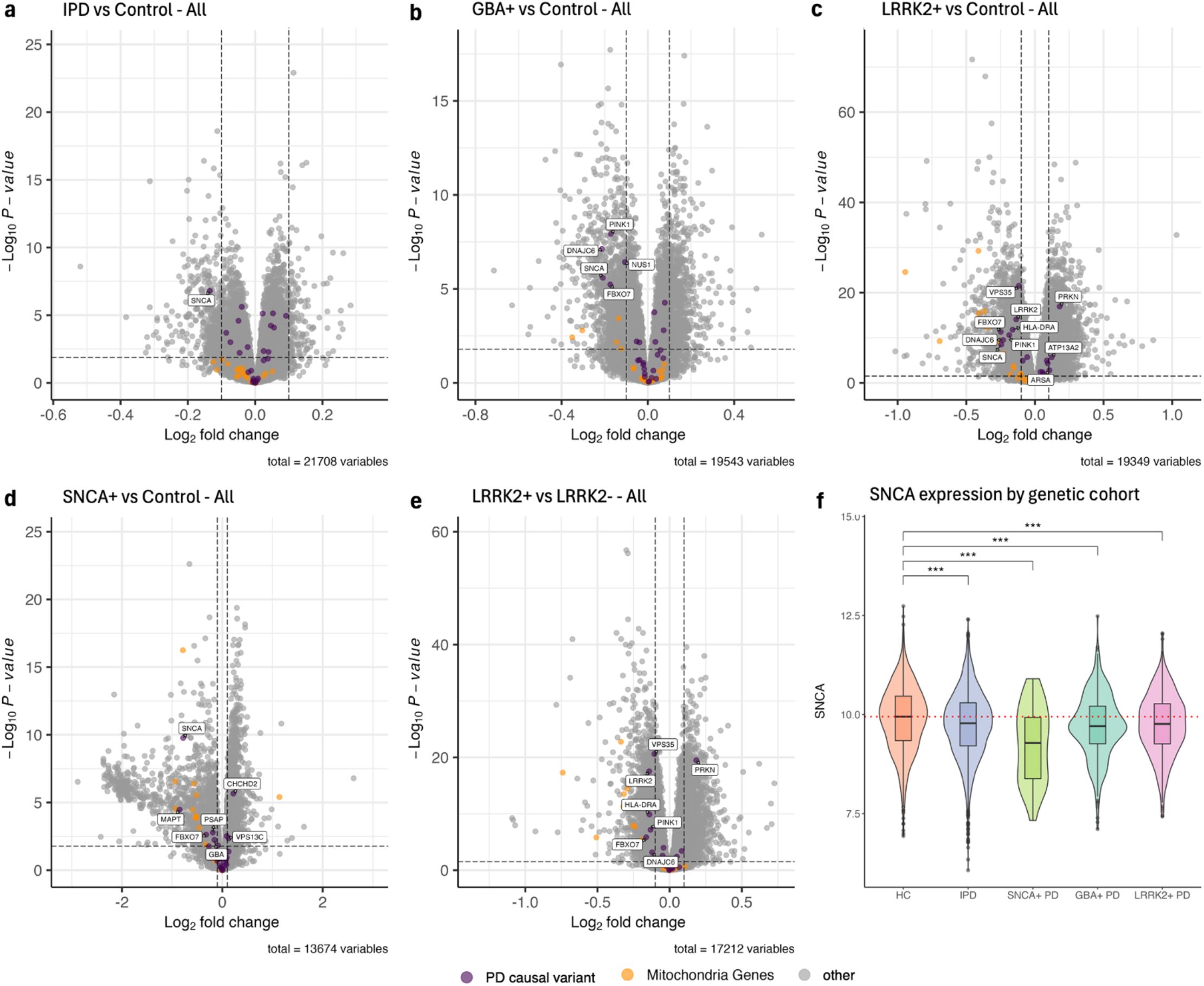
Differential expression analysis of all samples by genetic cohort. PD causal variants are colored purple, and mitochondrial genes are colored orange. Differentially expressed PD causal variants are additionally labeled by gene name. **a**, Idiopathic case samples with no SNCA/LRRK2/GBA mutation were compared to control samples with no PD-related mutations. **b**,**c**,**d**, Control samples with no PD-related mutations were compared to case samples with GBA+ (**b**), LRRK2+ (**c**), and SNCA+ (**d**) mutations. **e**, LRRK2+ case samples were compared to LRRK2-case samples. **f**, log(CPM) SNCA expression in HC, IPD, SNCA+ case, GBA+ case, and LRRK2+ case samples corrected by predicted neutrophil percentage, stratified by genetic cohort. HC includes control samples with no SNCA/LRRK2/GBA mutations. Adjusted p-values are labeled according to the adjusted p-value of SNCA differential expression from respective DE analyses. The dotted red line represents the median SNCA expression in HC samples.

We then conducted DE analysis between specific PD cohort by genetic status and compared these samples to health control samples (where healthy control refers to participants who were labeled ‘Control’ at baseline and have no SNCA+/GBA+/LRRK2+ mutation). We continue to see a statistically significant downregulation of SNCA in each PD cohort vs HC analysis, with a similar improvement in SNCA DE signal compared to uncorrected analysis (Fig. 4a,b,c,d, Supplemental Fig. 2). SNCA is not differentially expressed in the LRRK2+ vs LRRK2-comparison, which is consistent with our finding that SNCA is only downregulated in relation to control samples (Fig. 4e). The number of neutrophil- and lymphocyte-enriched genes were also not meaningfully decreased in LRRK2+ vs LRRK2-after correcting for predicted neutrophil percentage (Supplemental Fig. 6f,l), suggesting a stronger relationship between LRRK2 and neutrophils in whole blood consistent with the known interactions between LRRK2 and neutrophils.

### Pathway analysis highlights mitochondrial dysfunction in PD

To identify pathways with altered expression in the PD cohorts, we conducted Ingenuity Pathway Analysis (IPA) on differential expression results from each PD vs control cohort analysis. As found in our prior work, the most significantly enriched pathway in PD cases vs controls without correcting for neutrophil percentage was ‘Neutrophil Degranulation’ (Supplemental Fig. 7). IPA of neutrophil-corrected DE results successfully eliminates the ‘Neutrophil Degranulation’ pathway, with the most significantly enriched pathway now being ‘Mitochondrial RNA degradation’ (Supplemental Fig. 8). Neutrophil correction overall reduces the number of pathways with z-scores greater than 2 or less than -2 and p-values less than 0.05 from 45 without correction to 12 with. Notably, the ‘Parkinson’s Signaling Pathway’ becomes significantly expressed in the PD samples with neutrophil correction (Supplemental Figures).

When conducting pathway analysis by genetic cohort, even with neutrophil-corrected DE analysis, IPA still indicates significant enrichment of blood cell and immune-related pathways. As gene expression data was sequenced from blood samples, high enrichment of these pathways can likely be attributed to tissue type. Considering this possibility, we instead focused on pathways related to potentially relevant aspects of PD identified in prior studies, especially mitochondrial-related activity due to the significant presence of such pathways in the neutrophil-corrected case vs control IPA results. Pathways related to mitochondrial function were significantly enriched in every PD cohort vs control analysis (Supplemental Fig. 8-12). We particularly see positive enrichment of the ‘Mitochondrial Dysfunction’ pathway in GBA+, SNCA+, and LRRK2+ samples. Additionally, other pathways such as ‘Granzyme A Signaling’ in LRRK2+, ‘NAFLD Signaling Pathway’ in SNCA+, and ‘Coronavirus Pathogenesis Pathway’ in GBA+ are positively enriched and involved in mitochondrial dysfunction. In the IPD vs HC analysis, ‘BBSome Signaling Pathway’, a pathway positively involved in mitochondrial function, was one of nine significantly enriched pathways and was found to be depressed in IPD samples^8^. ‘Leukocyte Extravasation Signaling’ was another mitochondria-related pathway positively significant in the IPD analysis, involved in mitochondrial fission and fusion. This association between PD and mitochondrial activity is further supported by differential expression analysis, as genes in the mitochondrial genome are consistently downregulated in the PD cases vs controls comparisons (Fig. 3b,d). Mitochondrial gene downregulation appears more evident after neutrophil correction (Fig. 3d). We again see a similar overall downregulation of mitochondrial genes in DE analyses split by PD cohort (Fig. 4a,b,d,c). LRRK2+ samples appear to have both lower mitochondrial gene expression and a larger number of significantly enriched mitochondria-related pathways compared to all other PD cohorts (Fig. 4e, Supplemental Fig. 12). We can overall see consistent evidence of mitochondrial activity in relation to PD, with a specific trend towards increased dysfunction in disease samples.

### PD cohorts exhibit niches of gene expression in unsupervised analysis

The presence of differentially expressed genes and enriched pathways from DE and IPA analyses suggest different cohorts PD cohorts may exhibit different overarching transcriptomic profiles. To further investigate any global transcriptomic differences between PD and control cohorts, we applied UMAP dimensionality reduction and tried to identify any clusters of HC, IPD, SNCA+ PD, GBA+ PD, and LRRK2+ PD samples. From pathway analysis, we established an enrichment of mitochondrial pathways in PD whole blood gene expression, especially related to mitochondrial dysfunction. As such, we performed UMAP dimensionality reduction on genes in the Parkinson’s Signaling Pathway (Fig. 5a), the Mitochondrial Dysfunction pathway (Fig. 5b), BBSome Signaling Pathway (Fig. 5c), and Leukocyte Extravasation Signaling (Fig. 5d), as listed in the IPA database.

**Fig. 5.**
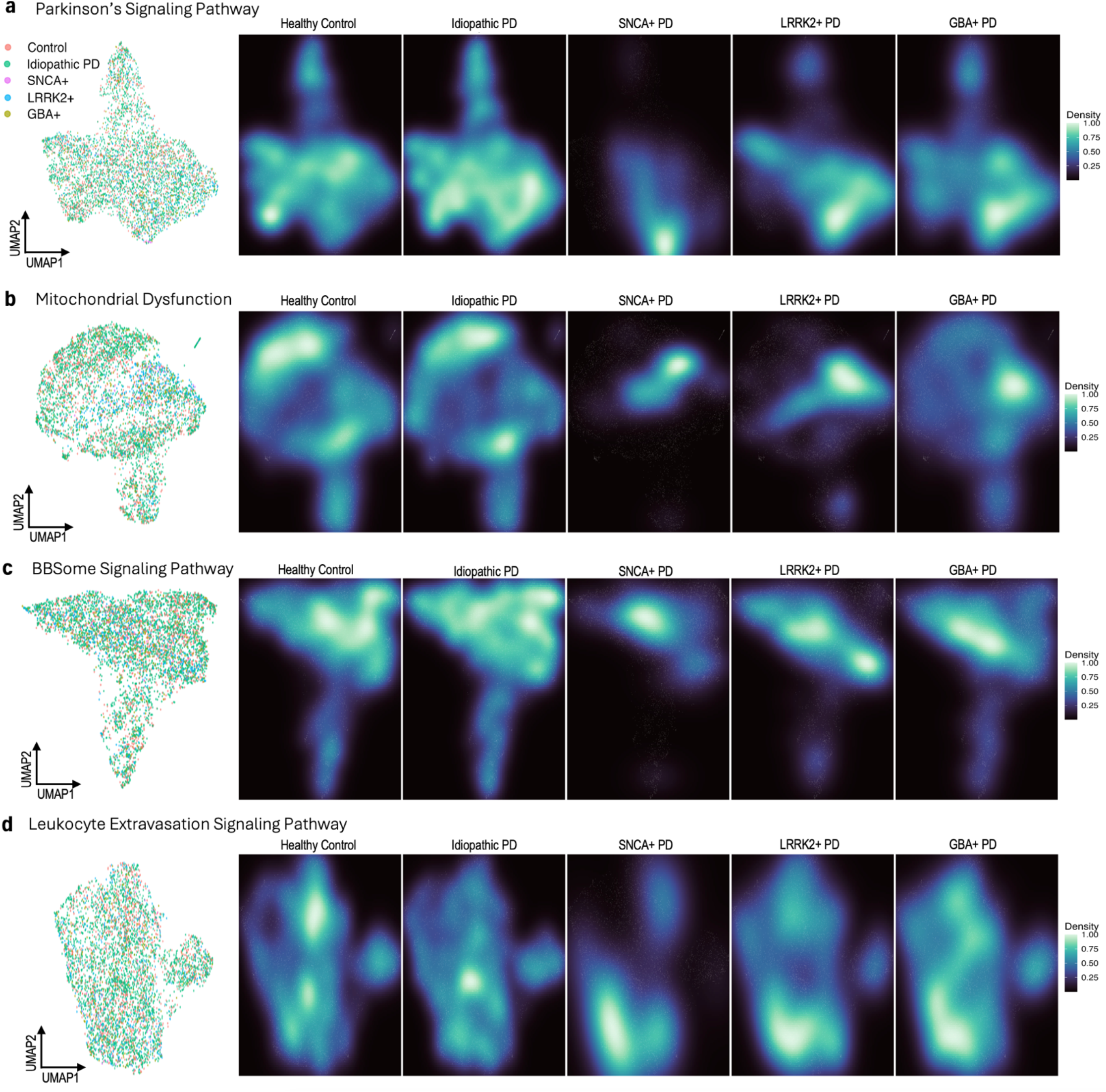
UMAP dimensionality reduction with pathway-specific genes by disease status and genetic cohort. UMAP embeddings were created from variant stabilizing transformed (VST) gene counts. Counts were then corrected for participant age and sex, as well as sample mRNA percentage and predicted neutrophil percentage. **a**, UMAP of 36 genes found in the Parkinson’s Signaling Pathway, with samples labeled by disease and genetic status, excluding samples from participants with unknown genetic status (n = 5,470). Corresponding density plots were made, stratified by disease and genetic status. **b**, A UMAP and set of density plots were created from 143 genes in the Mitochondrial Dysfunction pathway (n = 5,339). A small cluster of samples, mostly Control and Idiopathic PD, were removed from the Mitochondrial Dysfunction UMAP for visualization purposes. **c**, A UMAP of 59 BBSome Signaling Pathway genes and density plots (n = 5,358). A small cluster of samples, mostly Control and Idiopathic PD, were removed for visualization purposes. **d**, UMAP and density plots of 52 Leukocyte Extravasation Signaling Pathway genes (n = 5,345). A small cluster of samples, mostly Control and Idiopathic PD, were removed for visualization purposes.

None of the four UMAPs exhibit obvious clustering by cohort, even between control and case samples. Seeing as hierarchal clustering also failed to discriminate between cohorts, we can infer that PD whole blood gene expression may have no unique global transcriptomic structure compared to healthy controls, even when considering only disease-specific pathways (Supplemental Figure 13). However, by creating density plots of each UMAP stratified by sample cohort, we can begin to see specific niches of gene expression, especially within SNCA+, GBA+, and LRRK2+ samples. In the UMAP of genes in the Parkinson’s Signaling Pathway, we can see that while HC and IPD samples are present at a relatively consistent level across the entire plot, genetic cohorts tend to cluster near the bottom right (Fig. 5c). In mitochondrial dysfunction genes, we can see an even more distinct division between HC/IPD and SNCA+/GBA+/LRRK+, where genetic samples predominantly cluster in a central band and HC/ID samples cluster more towards the top and bottom of the UMAP (Fig. 5d). SNCA+ samples tend to occupy the smallest niche of expression in both Parkinson’s signaling pathway and mitochondrial dysfunction genes. Differences between IPD and HC samples, however, are comparatively minor. Some minute differences can still be observed, including in the UMAP density plots based on the IPD-enriched BBSome Signaling Pathway and Leukocyte Extravasation Signaling (Fig. 5c,d).

### Demographic, clinical, and biological factors influencing SNCA gene expression

The most striking results from our analysis of PD whole blood RNA sequencing data has been the significant downregulation of SNCA in DE analyses. We further investigated which phenotypic characteristics of PD could be responsible for the SNCA signal by plotting CPM normalized and logged SNCA gene counts by mutation status and age at baseline.

Lower SNCA expression compared to HC is most apparent in SNCA+ samples, both in those from participants with a clinical PD diagnosis and without (Fig. 6a). This decreased expression is statistically significant even when only considering samples taken at baseline (Fig. 6b). Mutations in SNCA are typically missense, indicating that transcriptomic depression of the gene may be due to some indirect mechanism that potentially becomes active in PD^24^. Combined with the fact that SNCA-PD samples also exhibit SNCA downregulation, it is apparent that SNCA mutations alone are not responsible for the decreased expression. Looking at other potential genetic drivers, we do see a more modest decrease of SNCA in LRRK2+ and GBA+ PD samples, which is statistically significant when considering expression in all samples but becomes unsignificant in samples taken at baseline (Fig. 4f, Fig. 6a,b). This is likely due to the relatively smaller pool of LRRK2+ PD and GBA+ PD participants compared to HC and more subtle decrease in SNCA expression compared to SNCA+ PD participants (Fig. 6b). The overall behavior of SNCA expression is consistent with our findings in differential expression analysis by genetic cohort (Supplemental Fig. 3). It remains likely that LRRK2 and GBA, along with SNCA, may indirectly impact whatever mechanism is responsible for decreased SNCA in PD.

**Fig. 6.**
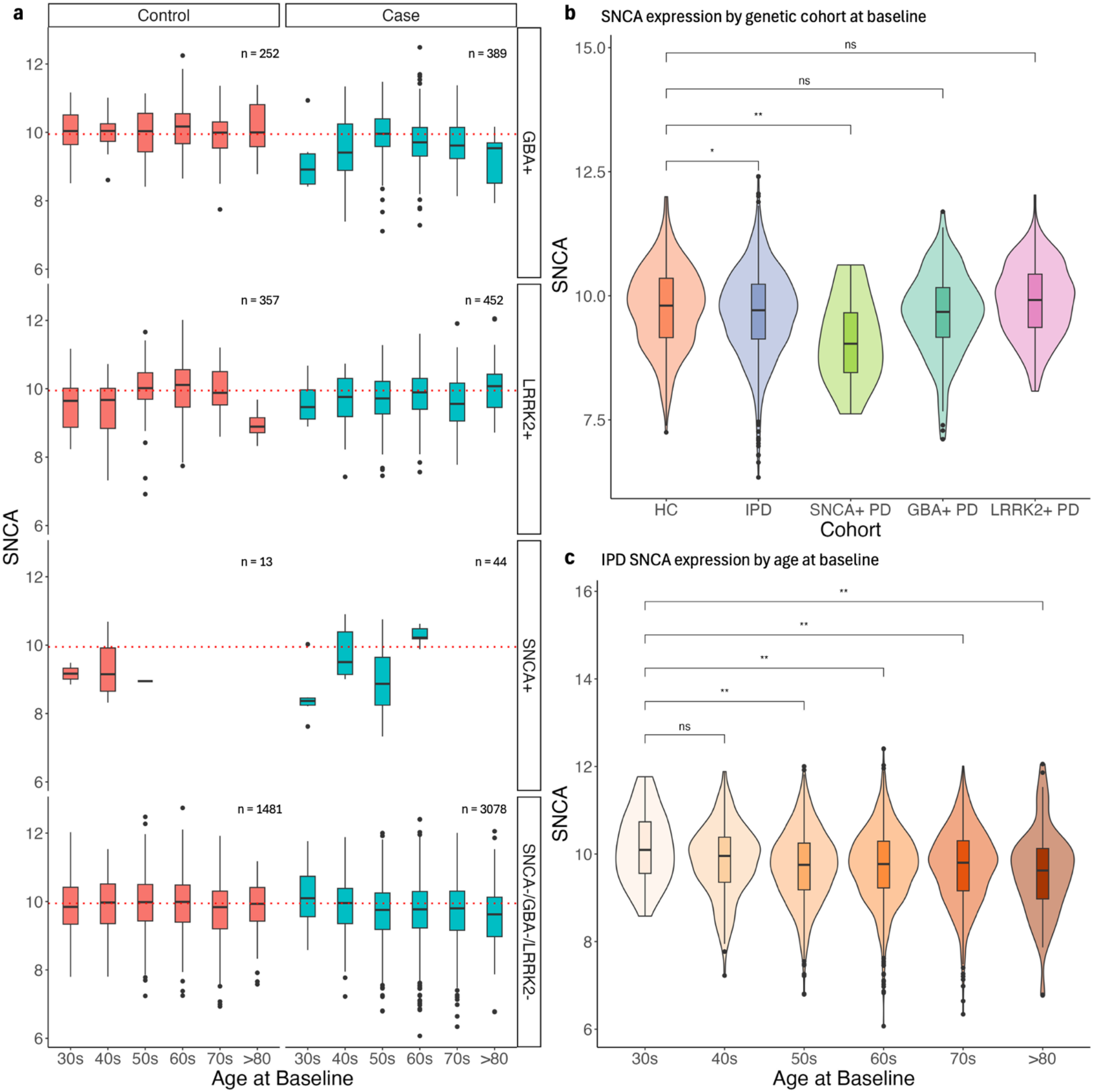
SNCA expression stratified by demographic, clinical, and biological factors. **a, b, c**, Gene counts were log(CPM) normalized and corrected for predicted neutrophil percentage. **a**, SNCA expression in samples stratified by genetic status and diagnosis. The red dotted line represents the median SNCA expression in healthy control samples (i.e. ‘Control’ and ‘SNCA-/GBA-/LRRK2-’). **b**, SNCA expression of HC, IPD, SNCA+ case samples, GBA+ case samples, and LRRK2+ case samples at baseline only. **c**, SNCA expression of IPD samples over age at baseline. All p-values were calculated using a Wilcoxon rank-sum test.

Further analysis of SNCA expression by participant age at baseline also suggests a possible SNCA-age dependency in PD. In IPD samples (i.e. Case and SNCA-/GBA-/LRRK2-), there is a statistically significant decrease in SNCA expression as the baseline age of the participant increases (Fig. 6c). This steady age-related decrease of SNCA is not present in the control samples of any genetic cohort, indicating a possible disease-specific behavior of SNCA expression in whole blood. Overall, it appears that some combination of genetic and age-related factors may contribute to a systemic downregulation of SNCA in the PD whole blood transcriptome.

## Discussion

The goal of this study was to investigate the impact of blood cell-enriched gene expression on differential expression analysis of PD whole blood RNA sequencing samples. To correct for a previously established enrichment of neutrophil-enriched genes and the neutrophil degranulation pathway in PD case vs control DE and IPA analysis, we developed a linear model using 1,254 PPMI samples with CBC data to predict neutrophil percentage in 5,643 PPMI and PDBP samples. We developed four models: a linear model using blood cell-enriched gene expression, a second linear model with genes selected through MI feature selection, a third linear model using a combined set of the most significant genes from the prior two models, and a XGBoost regression model trained on the entire gene expression dataset. We selected the combined model for neutrophil prediction due to strong performance in the R-squared, RMSE, and MAE evaluation.

The combined 1,254 known and 5,643 predicted neutrophil percentages were then included as a design covariate in differential expression analysis, which successfully eliminated a large portion of neutrophil-enriched genes, as well as genes enriched in other blood cells, such as lymphocytes, monocytes, basophils, eosinophils, and dendritic cells. IPA analysis of neutrophil-corrected DE results no longer exhibit enrichment of the Neutrophil Degranulation pathway, indicating that correction using the predicted neutrophil percentages adequately accounts for pathway activity related to neutrophil function.

With neutrophil-correction, we continue to see a consistent depression and DE signal improvement of SNCA expression in the whole blood transcriptome for all PD cohorts. Why we see SNCA downregulation is an open question. For one, SNCA appears to be relevant to PD beyond the gene’s involvement in blood cell function. While SNCA is not highly expressed in neutrophils, the gene is expressed in plasmacytoid dendritic cells (115.5 pTPM), classical monocytes (62.7 pTPM), and basophils (36.8 pTPM)^20^. We demonstrated that neutrophil correction decreases the number of differentially expressed blood cell-enriched genes across all cell types, which may explain the greater magnitude and significance of SNCA downregulation after correction. Including neutrophil percentage in our analyses appears to uncover a stronger gene expression signal for SNCA in PD cohorts that occurs independently of immune cells. The consistent depression of SNCA in every PD cohort compared to healthy controls, including in IPD samples without SNCA+/GBA+/LRRK+, suggests that SNCA downregulation in whole blood transcriptomic analysis may be a notable hallmark of disease.

We further identified multiple mitochondria-related pathways enriched in PD cohorts with IPA. Most notably is the positive enrichment of the Mitochondrial Dysfunction pathway in SNCA+, GBA+, and LRRK2+ samples. SNCA is known to be involved in mitochondrial function and is present in the Mitochondrial Dysfunction pathway specifically, suggesting that mitochondrial activity may be what is driving the SNCA DE signal in PD. Overly active mitochondria has previous been tied to neuronal cell death and neurogenerative disease^25^. Mitochondrial inhibition in DA neurons was also demonstrated in multiple studies to induce parkinsonian motor symptoms in both primates and humans^26,27^. Barnhoorn et al. similarly found a reduction of mitochondrial function in PD within the PPMI samples used in this study, and that mitochondrial dysfunction appears to scale with disease severity in gene set enrichment analysis^28^. We see further evidence of mitochondrial activity differences in UMAP density plots, where PD cohorts exhibit tendencies to cluster toward specific niches of gene expression rather than a fully unique transcriptomic profile. The lack of distinct global structural differences between PD cohorts and control samples corresponds with the high degree of complexity in PD development and expression. It is possible that more sophisticated methods of unsupervised analysis may provide a better understanding of the highly nuanced transcriptomic differences in PD related to mitochondrial activity and dysfunction.

This study does have notable limitations. For one, despite utilizing the largest compiled PD transcriptomic dataset, the analyses in this study were limited to clinical diagnosed PD participants and control individuals due to the relatively low number of prodromal and SWEDD patients. We also conducted analyses by sample rather than by participant, since individuals with SNCA/GBA/LRRK2 mutations, especially SNCA, compose a relatively small proportion of our total pool of participants. As such, participant-related sample dependencies may be unaccounted for in some statistical tests. While SNCA may be a strong signal of PD, significant transcriptomic differences that typify IPD are still unclear, and will likely require further single-cell level analysis to elucidate. Overall, future work in single-cell transcriptomic analysis of blood cells will be necessary to identify which differentially expressed genes and pathways can be pathogenically linked to PD, and which are merely associated with the disease.

In conclusion, this study incorporated neutrophil percentage correction into differential transcriptomic analysis of PD whole blood samples. We see a stronger depression of SNCA expression, which may be caused by mitochondrial dysfunction and other related mitochondrial pathway activity. Correcting for immune cell-enriched genes in PD whole blood RNA analysis can uncover more relevant pathways in the PD transcriptomic profile, which will help guide future work in blood-based analysis of the disease.

## Methods

### Data

All data used in the study was collected and processed according to the protocols outlined by the Parkinson’s Progression Markers Initiative (PPMI) and Accelerating Medicines Partnership Parkinson’s Disease (AMP PD®) program^11,12^. Both PPMI and Parkinson’s Disease Biomarkers Program (PDBP) from AMP PD® patient samples were labeled by their disease status (PD or non-PD) and genetic group (GBA+/-, LRRK2+/-, and SNCA+/-mutation status). A subset of PPMI patient samples included complete blood count (CBC) and neutrophil percentage.

### Genome and transcriptome alignment, quantification and quality control

Samples were sequenced, aligned, and quantified as part of PPMI. Sequencing was done using the Illumina NovaSeq 6000 platform, after which FASTQ files were aligned to the GRCh38 human genome using STAR 2.6.1d. Gene counts were created using featureCounts 1.6.2 and GENCODE 29 annotations.

A total of 8,461 samples were provided by PPMI and AMP PD® (labeled as either PDBP or BioFIND). Only samples with a RIN value of greater than 6, usable bases percentage between 20% and 65%, and chimeric reads percentage below 3% were included in both neutrophil percentage prediction model development and differential gene expression analysis. Additionally, 166 BioFIND samples were removed due to the relatively small sample size compared to PPMI and PDBP. Of the 6,490 passing samples, 1,254 were used for neutrophil prediction model development. 6,490 samples from participants labeled as ‘Case’ or ‘Control’ at baseline were included in differential expression analyses.

### Neutrophil percentage linear modeling and prediction

Genes enriched in white blood cells (neutrophils, eosinophils, basophils, monocytes, lymphocytes, and dendritic cells) were identified using annotations from the Human Blood Atlas^20^. Linear models were developed using the lm() function in R. Backward elimination was applied recursively until the only genes used in the model had p-values less than 0.05. Mutual information features selection was conducted using SelectKBest() and mutual_info_regression from sklearn in python. The XGBoost regression model was built using the xgboost r package with the parameters nrounds = 10, eta = 0.3 and max depth = 3. All four models were compared using the average Pearson R-squared, root mean squared error and mean absolute error across 100 train-test splits.

### Variance analysis

PCA was calculated using the plotPCA() function from DESeq2 and batch correction was conducted with removeBatchE5ect() function from limma. PCs were correlated to the technical (study, plate, usable bases, etc.) and biological (neutrophil percentage, age, sex, etc.) variables of the samples using a Spearman’s rank correlation or intraclass correlation if the variable were continuous or categorical respectively. The significance of each correlation was evaluated using either a spearman or ANOVA test p-value for continuous or categorical variables respectively. Categorical variables with singular unique values (sample_id, participant_id, diagnosis_at_baseline, diagnosis_latest) could not be evaluated using either method, and as such p-values were set to 0 for all PCs.

### Differential expression analysis

All differential expression analyses were conducted using the well-developed limma-voom 3.58.1 framework in R. We used a p-value threshold of 0.05 and log fold change threshold of 0.1. Each comparison used the design = ∼0 + case + sex + percent mRNA bases + predicted neutrophil percentage + age squared, where age is determined by the age at patient enrollment. The design matrix was determined through successive testing and variance analysis to identify the most significant and biologically relevant covariates (Supplemental Fig. 1, Supplemental Table 4).

Genes were labeled in volcano plots by category of interest. Causal variants of PD were determined by evidence in previous studies. Mitochondrial genes were labeled based on genes in the Human Gene Nomenclature Committee’s mitochondrial genome list^29^. Pathway genes were compiled and identified from statistically significant pathways in Ingenuity Pathway Analysis (IPA) analyses. Leukocyte-enriched genes were identified per the Human Blood Atlas.

### Ingenuity pathway analysis

Differential expression analysis results from multiple comparisons were used to conduct pathway analysis with QIAGEN Ingenuity Pathway Analysis (IPA) software (QIAGEN Inc., https://digitalinsights.qiagen.com/IPA). For IPA of DE results without neutrophil correction, an adjusted p-value threshold of 0.05 and log fold change threshold of < -0.1 and > 0.1 was applied to identify significant genes. For IPA of DE results with neutrophil correction, an adjusted p-value threshold of 0.05 and log fold change threshold of < -0.085 and > 0.065 was used to avoid biased z-score calculations and include a similar number of differentially expressed genes as the un-corrected analysis (∼500 upregulated and ∼500 downregulated). Log fold change thresholds of < -0.1 and > 0.075 and an adjusted p-value threshold of 0.05 were used for IPD vs HC analysis. GBA+ v HC analysis was conducted with the same adjusted p-value threshold and a log fold change threshold of < -0.1 and > 0.09. LRRK2+ v HC was conducted with the same adjusted p-value threshold and a log fold change threshold of < -0.14 and > 0.15. Finally, SNCA+ v HC was conducted with the same adjusted p-value threshold and a log fold change threshold of < -0.1 and > 0.14. Genes from significant pathways related to mitochondrial function were then compiled and used in UMAP dimensionality reduction.

### Dimensionality reduction and density plots

Uniform Manifold Approximation and Projection (UMAP) dimensionality reduction was conducted using the umap() function the R package umap version 0.2.10.0. Gene counts were normalized and transformed using DESeq2 vst() before applying dimensionality reduction. To make the density plots, samples were labeled with ‘Control’ if the participant was not diagnosed with PD and was SNCA-/GBA-/LRRK2-, ‘Idiopathic PD’, if the participant was diagnosed with PD and was SNCA-/GBA-/LRRK2, SNCA+ if the participant was diagnosed with PD and had a SNCA mutation, GBA+ if the participant was diagnosed with PD and had a GBA mutation, and LRRK2+ if the participant was diagnosed with PD and had a LRRK2 mutation. Density plots were created using ggplot2 and the stat_density_2d() function. Mitochondrial genes were identified using the HUGO Gene Nomenclature Committee (HGNC) Mitochondrial genome gene group. Mitochondrial dysfunction, BBSome signaling pathway, Leukocyte Extravasation Signaling, and Parkinson’s Signaling Pathway genes were identified based on the corresponding molecule list in IPA.

### Statistical software

Statistical analyses were conducted in either R version 4.3.1 or python version 3.7.16. Linear models were created in R using the lm() from the R stats package and the XGBoost model was built using the R package xgboost version 1.7.8.1. The package scikit-learn version 1.0.2 was used for mutual information feature selection in python. Differential expression analysis and variance analysis were conducted in R using DESeq2 version 1.40.2, limma version 3.58.1, and edgeR version 3.42.4. Plots were made using either ggplot2 version 3.4.4 or EnhancedVolcano version 1.18.0. The geom_signif() function in ggpubr version 0.6.0 was used whenever a two-tailed Wilcoxon rank sum test was applied.

## Data availability

Raw sequencing data (FASTQ files), alignment files (BAM files), TPM data and counts for each sample are available at the LONI IDA. (https://fairsharing.org/, IDA; LONI IDA, https://doi.org/10.25504/FAIRsharing.r4phfff). Data are also available through the AMP PD® (https://amp-pd.org/). These are the requirements for downloading from the AMP PD®: (1) personal and institutional or company details; (2) description of intended data use, for example, proposed analyses; (3) institutional signature on the AMP PD® Data Use Agreement (for researchers requesting access to individual level, ‘omics data). Additional data, including but not limited to study arm, motor assessments, DaTscan and MRI imaging, genetic testing results, whole-exome and genome sequencing data, patient history and standardized techniques and protocols for data collection are also available through the IDA. To access complete data, researchers need to fill out a data-use agreement. Data are available in a public (institutional, general or participant-specific) repository that does not issue datasets with DOIs (non-mandated deposition).

## Code availability

All code for data analysis in this study is available on GitHub and can be accessed via this link: https://github.com/kaylaxu/pd_wb_rnaseq_snca_paper.

## Acknowledgements

This research was supported in part by a grant from NINDS (U01-NS120260) (KX, DWC, and IV). This research was supported and funded by the MJFF under grant numbers 12749 (K.V.K.-J.), 12749.01 (K.V.K.-J., M.R.C.) and 14696 (K.V.K.-J., A.K., D.W.C.). Many MJFF sta5 assisted in harmonizing and transferring data. This research was supported in part by the Intramural Research Program of the National Institutes of Health, National Institute on Aging. Data used in the preparation of this article were obtained on 2022-12-09 from the Parkinson’s Progression Markers Initiative (PPMI) database (https://www.ppmi-info.org/access-data-specimens/download-data), RRID:SCR_006431. For up-to-date information on the study, visit http://www.ppmi-info.org. PPMI – a public-private partnership – is funded by the Michael J. Fox Foundation for Parkinson’s Research and funding partners, including 4D Pharma, Abbvie, AcureX, Allergan, Amathus Therapeutics, Aligning Science Across Parkinson’s, AskBio, Avid Radiopharmaceuticals, BIAL, BioArctic, Biogen, Biohaven, BioLegend, BlueRock Therapeutics, Bristol-Myers Squibb, Calico Labs, Capsida Biotherapeutics, Celgene, Cerevel Therapeutics, Coave Therapeutics, DaCapo Brainscience, Denali, Edmond J. Safra Foundation, Eli Lilly, Gain Therapeutics, GE HealthCare, Genentech, GSK, Golub Capital, Handl Therapeutics, Insitro, Jazz Pharmaceuticals, Johnson & Johnson Innovative Medicine, Lundbeck, Merck, Meso Scale Discovery, Mission Therapeutics, Neurocrine Biosciences, Neuron23, Neuropore, Pfizer, Piramal, Prevail Therapeutics, Roche, Sanofi, Servier, Sun Pharma Advanced Research Company, Takeda, Teva, UCB, Vanqua Bio, Verily, Voyager Therapeutics, the Weston Family Foundation and Yumanity Therapeutics. We also thank the AMP PD® for allowing us access to PDBP data. The PDBP consortium is supported by the National Institute of Neurological Disorders and Stroke at the National Institutes of Health. A full list of PDBP investigators can be found at https://pdbp.ninds.nih.gov/policy. PDBP investigators have not participated in reviewing the data analysis or content of the text. Data used in the preparation of this article were obtained from the AMP PD® Knowledge Platform. For up-to-date information on the study, visit https://www.amp-pd.org. The AMP PD®, a public–private partnership, is managed by the FNIH and funded by Celgene, GlaxoSmithKline, the MJFF, the National Institute of Neurological Disorders and Stroke, Pfizer, Sanofi and Verily. We thank all people with PD and families for participating in the study and donating their samples and time. We would like to thank the group at IU for RNA isolation, QC metrics and safe shipment of samples.

---

**Table 1.**
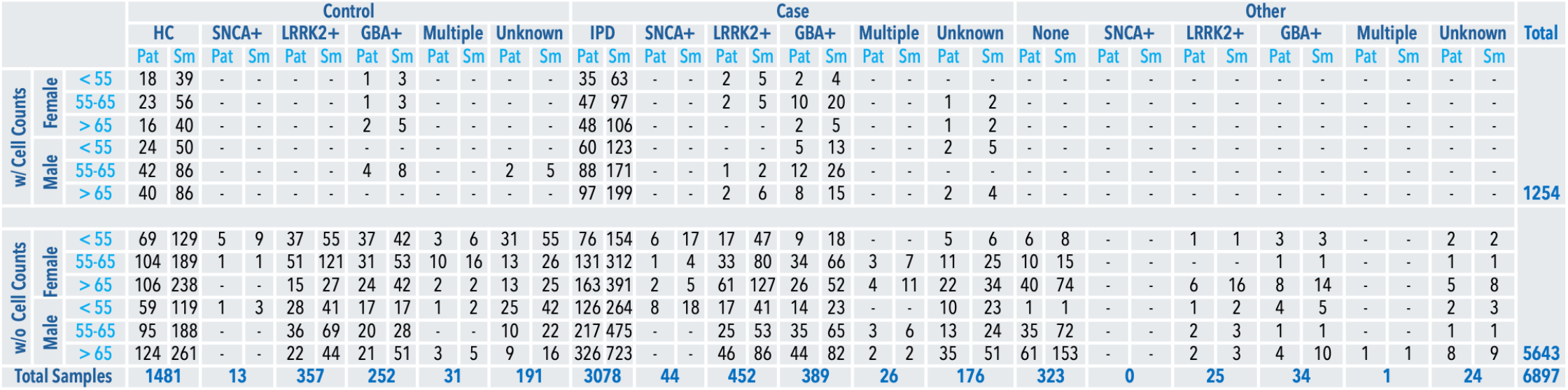
Demographic information and genetic status of PPMI and PDBP participants (Pat) and samples (Sm).

