## Supplemental Figures for "Decreased SNCA Expression in Whole-Blood RNA Analysis of Parkinson’s Disease Adjusting for Lymphocytes"

| | Coefficient | Std Error | T value | P value |
|---|---|---|---|---|
| **(Intercept)** | -166.3 | 22.40 | -7.423 | 0 |
| **ENSG00000205927** | -0.5728 | 0.2411 | -2.376 | 0.01771 |
| **ENSG00000113389** | 3.550 | 0.6763 | 5.249 | 0 |
| **ENSG00000164047** | 1.194 | 0.2233 | 5.346 | 0 |
| **ENSG00000158497** | -1.994 | 0.9674 | -2.061 | 0.03955 |
| **ENSG00000061273** | 1.727 | 0.4345 | 3.976 | 0 |
| **ENSG00000167186** | -2.563 | 0.9489 | -2.701 | 0.007039 |
| **ENSG00000151012** | 1.983 | 0.4868 | 4.074 | 0 |
| **ENSG00000133687** | -0.4661 | 0.1285 | -3.627 | 0.0003017 |
| **ENSG00000134827** | 1.403 | 0.4792 | 2.928 | 0.003487 |
| **ENSG00000172292** | -2.956 | 0.5767 | -5.126 | 0 |
| **ENSG00000038219** | 5.337 | 0.6258 | 8.529 | 0 |
| **ENSG00000158488** | -2.556 | 0.6875 | -3.718 | 0.0002124 |
| **ENSG00000135373** | 2.603 | 0.8006 | 3.252 | 0.001186 |
| **ENSG00000204345** | -0.8332 | 0.2993 | -2.784 | 0.005477 |
| **ENSG00000062716** | 8.633 | 0.6668 | 12.95 | 0 |
| **ENSG00000050748** | -4.712 | 0.9498 | -4.961 | 0 |
| **ENSG00000164187** | -2.656 | 0.5261 | -5.047 | 0 |
| **ENSG00000124721** | 1.668 | 0.4876 | 3.421 | 0.0006495 |
| **ENSG00000108846** | -0.6706 | 0.2472 | -2.712 | 0.006796 |
| **ENSG00000118292** | 1.991 | 0.8279 | 2.405 | 0.01634 |
| **ENSG00000150051** | -2.048 | 0.7065 | -2.898 | 0.003836 |
| **ENSG00000149516** | -1.692 | 0.3243 | -5.219 | 0 |
| **ENSG00000120949** | 1.723 | 0.5864 | 2.938 | 0.003386 |
| **ENSG00000163162** | 4.798 | 0.7300 | 6.572 | 0 |
| **ENSG00000175793** | 3.463 | 0.8893 | 3.894 | 0.0001052 |
| **ENSG00000083290** | 2.315 | 0.8190 | 2.827 | 0.004799 |
| **ENSG00000092969** | -1.899 | 0.7666 | -2.477 | 0.01342 |

**Supplemental Table 1 | Blood cell-based linear model genes and coefficient estimates.**

| | Coefficient | Std Error | T value | P value |
|---|---|---|---|---|
| **(Intercept)** | -157.6 | 6.996 | -22.52 | 0 |
| **ENSG00000151948** | 2.138 | 0.7743 | 2.761 | 0.005869 |
| **ENSG00000153179** | -2.558 | 0.8106 | -3.156 | 0.001650 |
| **ENSG00000179299** | 1.688 | 0.4408 | 3.829 | 0.0001366 |
| **ENSG00000143226** | -4.342 | 0.7855 | -5.527 | 0 |
| **ENSG00000173110** | 1.362 | 0.4507 | 3.021 | 0.002583 |
| **ENSG00000196549** | -2.219 | 0.4847 | -4.579 | 0 |
| **ENSG00000062716** | 5.451 | 0.7661 | 7.115 | 0 |
| **ENSG00000146592** | -1.169 | 0.4500 | -2.598 | 0.009522 |
| **ENSG00000163162** | 6.500 | 1.043 | 6.234 | 0 |
| **ENSG00000102010** | 2.596 | 0.4706 | 5.521 | 0 |
| **ENSG00000181274** | -2.015 | 0.5764 | -3.495 | 0.0004975 |
| **ENSG00000138463** | -2.471 | 0.6754 | -3.659 | 0.00026661 |
| **ENSG00000148572** | 3.838 | 0.6504 | 5.902 | 0 |
| **ENSG00000197208** | 3.137 | 0.5838 | 5.373 | 0 |
| **ENSG00000128594** | 3.248 | 0.6099 | 5.326 | 0 |
| **ENSG00000070731** | 2.087 | 0.4887 | 4.271 | 0 |
| **ENSG00000165046** | 4.371 | 0.6230 | 7.016 | 0 |

**Supplemental Table 2 | Mutual information feature selection-based linear model genes and coefficient estimates.**

| | Coefficient | Std Error | T value | P value |
|---|---|---|---|---|
| **(Intercept)** | -119.1 | 19.45 | -6.124 | 0 |
| **ENSG00000205927** | -0.5725 | 0.2368 | -2.417 | 0.01583 |
| **ENSG00000113389** | 3.602 | 0.6545 | 5.502 | 0 |
| **ENSG00000164047** | 1.002 | 0.2224 | 4.507 | 0 |
| **ENSG00000167186** | -3.047 | 0.9139 | -3.335 | 0.0008862 |
| **ENSG00000151012** | 1.194 | 0.4800 | 2.488 | 0.01301 |
| **ENSG00000179299** | 1.429 | 0.3890 | 3.673 | 0.0002529 |
| **ENSG00000143226** | -3.080 | 0.6326 | -4.868 | 0 |
| **ENSG00000133687** | -0.3179 | 0.1252 | -2.539 | 0.01128 |
| **ENSG00000134827** | 1.067 | 0.4715 | 2.264 | 0.02381 |
| **ENSG00000172292** | -1.330 | 0.5919 | -2.248 | 0.02482 |
| **ENSG00000038219** | 3.638 | 0.7248 | 5.020 | 0 |
| **ENSG00000158488** | -2.470 | 0.6610 | -3.736 | 0.0001977 |
| **ENSG00000135373** | 2.594 | 0.7782 | 3.333 | 0.0008923 |
| **ENSG00000204345** | -0.7559 | 0.2935 | -2.576 | 0.0101458 |
| **ENSG00000196549** | -1.345 | 0.4162 | -3.232 | 0.001272 |
| **ENSG00000062716** | 6.168 | 0.7486 | 8.239 | 0 |
| **ENSG00000050748** | -5.986 | 0.9333 | -6.412 | 0 |
| **ENSG00000164187** | -2.387 | 0.5955 | -4.008 | 0 |
| **ENSG00000124721** | 1.213 | 0.4704 | 2.579 | 0.01005 |
| **ENSG00000108846** | -0.5261 | 0.2389 | -2.203 | 0.02785 |
| **ENSG00000150051** | -2.078 | 0.6796 | -3.058 | 0.002289 |
| **ENSG00000149516** | -1.494 | 0.3243 | -4.607 | 0 |
| **ENSG00000163162** | 4.403 | 0.9119 | 4.829 | 0 |
| **ENSG00000175793** | 3.267 | 0.8585 | 3.805 | 0.0001506 |
| **ENSG00000083290** | 3.250 | 0.7634 | 4.257 | 0 |
| **ENSG00000092969** | -2.166 | 0.7468 | -2.900 | 0.003810 |
| **ENSG00000148572** | 3.133 | 0.6347 | 4.937 | 0 |
| **ENSG00000197208** | 2.177 | 0.5319 | 4.092 | 0 |
| **ENSG00000128594** | 1.819 | 0.5224 | 3.483 | 0.0005188 |
| **ENSG00000070731** | 1.389 | 0.4644 | 2.992 | 0.002846 |
| **ENSG00000165046** | 1.571 | 0.5831 | 2.695 | 0.007159 |

**Supplemental Table 3 | Combined feature selection linear model genes and coefficient estimates.**

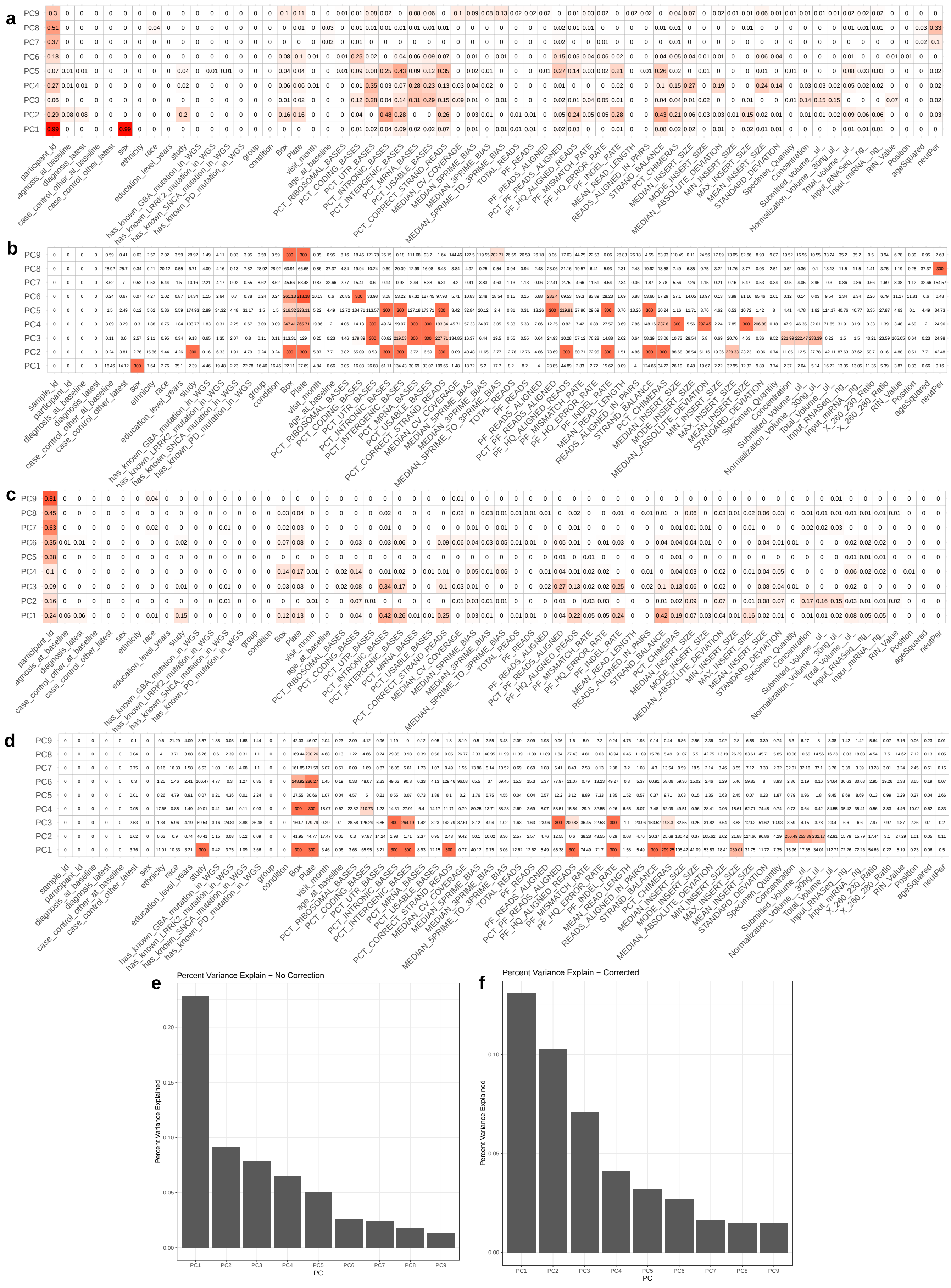


**Supplemental Figure 1 | PCA analysis of whole blood transcriptomic variance. a,b,c,d,** PCs were correlated to sample demographic, QC, and clinical data. Correlation with categorical values (such as participant_id, study, etc.) were calculated using the intraclass correlation coefficient. Continuous value correlations were calculated with spearman rho. Plots were created both without batch correction (**a**) with associated p-values (**b**) and with case, sex, predicted neutrophil percentage (neutPer), mrna base percentage, and age-squared correction (**c**) with associated p-values (**d**). All correlations are statistically significant. **e,f,** Plots of percent variance explained by each PC without correction (**e**) and with correction (**f**).

| Design | Upregulated | Downregulated | Total | SNCA Adjusted P-value rank |
|---|---|---|---|---|
| ~0 + case | 1772 | 1000 | 2772 | 1458 |
| ~0 + case + sex | 1523 | 765 | 2288 | 1320 |
| ~0 + case + sex + neutPer | 184 | 441 | 625 | 31 |
| ~0 + case + sex + neutPer + plate | 174 | 390 | 564 | 54 |
| ~0 + case + sex + neutPer + study | 217 | 384 | 601 | 38 |
| ~0 + case + sex + neutPer + ageSquared | 135 | 329 | 464 | 20 |
| ~0 + case + sex + neutPer + ageSquared + intronicBases | 126 | 334 | 460 | 21 |
| ~0 + case + sex + neutPer + ageSquared + mrnaBases | 110 | 357 | 467 | 18 |
| ~0 + case + sex + ageSquared + mrnaBases | 1224 | 530 | 1754 | 973 |

**Supplemental Table 4 | Design testing for PD differential expression analysis.** Testing was conducted with PD case vs all control samples. The design with the highest SNCA rank was used for all DE analyses in the study.

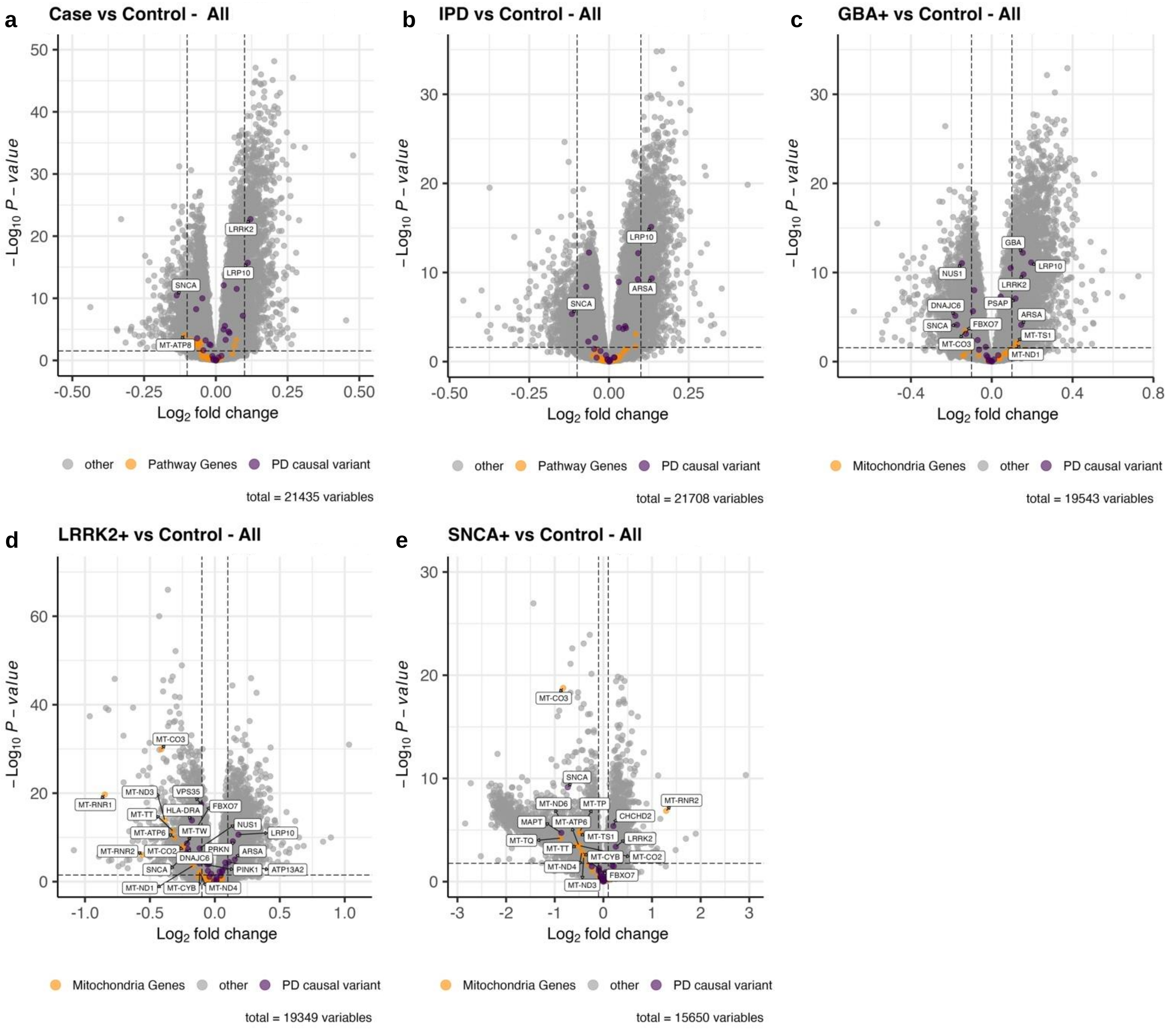


**Supplemental Figure 2 | Differential expression analysis of PD case cohorts vs control without predicted neutrophil percentage correction.** PD causal variants are colored purple, and mitochondrial genes are colored orange. Differentially expressed PD causal variants are additionally labeled by gene name. **a,** All PD cases were compared to all control samples based on the diagnosis at baseline. **b,** Idiopathic case samples with no SNCA+/LRRK2+/GBA+ mutation were compared to control samples with no PD-related mutations. **c,d,e,** Control samples with no PD-related mutations were compared to case samples with GBA+ (**c**), LRRK2+ (**d**), and SNCA+ (**e**) mutations.

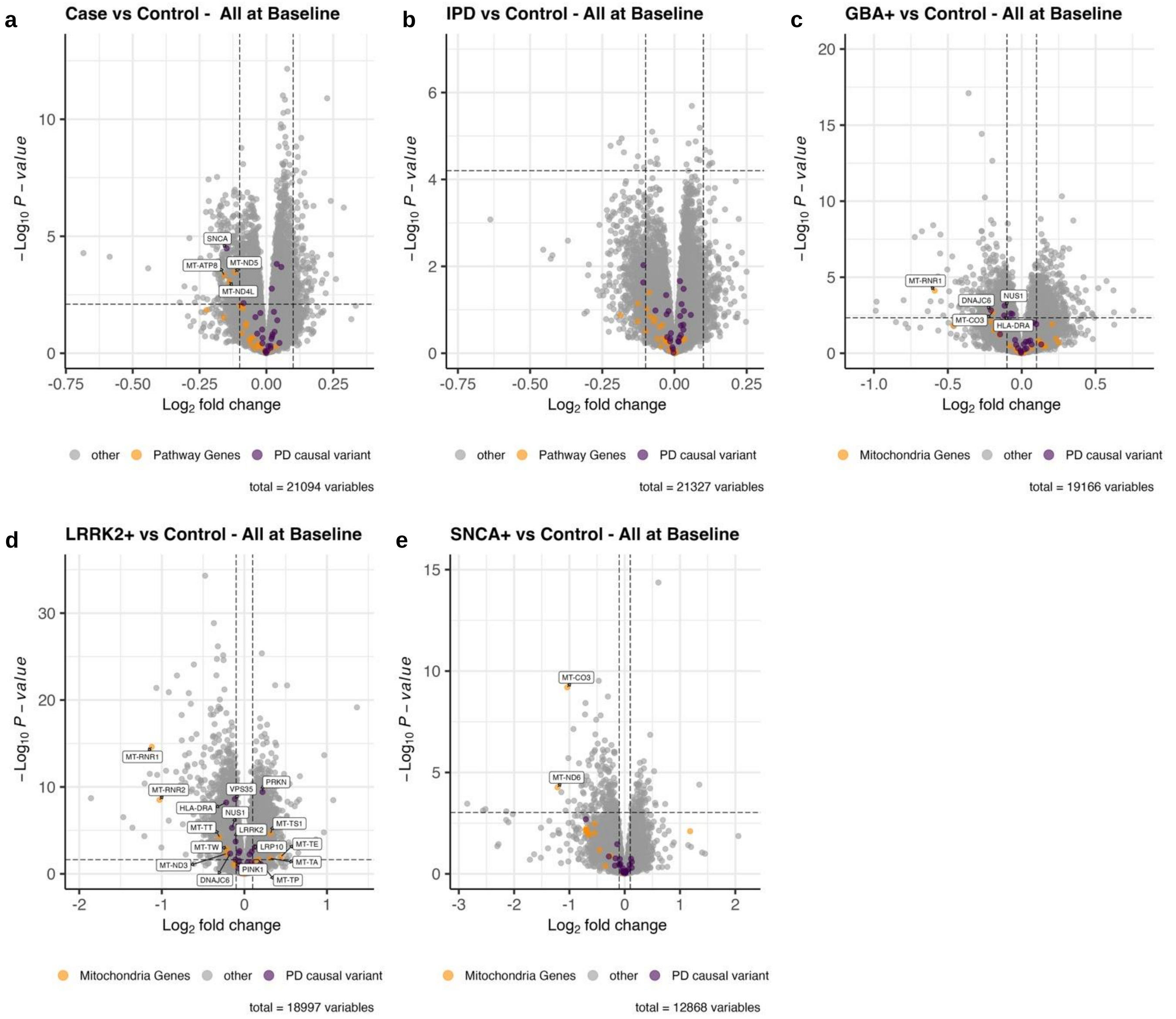


**Supplemental Figure 3. Differential expression analysis of PD case cohorts vs control at baseline with predicted neutrophil percentage correction.** All DE analyses were conducted with only baseline samples taken upon initial enrollment into either the PPMI or PDBP study. PD causal variants are colored purple, and mitochondrial genes are colored orange. Differentially expressed PD causal variants are additionally labeled by gene name. **a,** All PD cases were compared to all control samples based on the diagnosis at baseline. **b,** Idiopathic case samples with no SNCA+/LRRK2+/GBA+ mutation were compared to control samples with no PD-related mutations. **c,d,e,** Control samples with no PD-related mutations were compared to case samples with GBA+ (**c**), LRRK2+ (**d**), and SNCA+ (**e**) mutations.

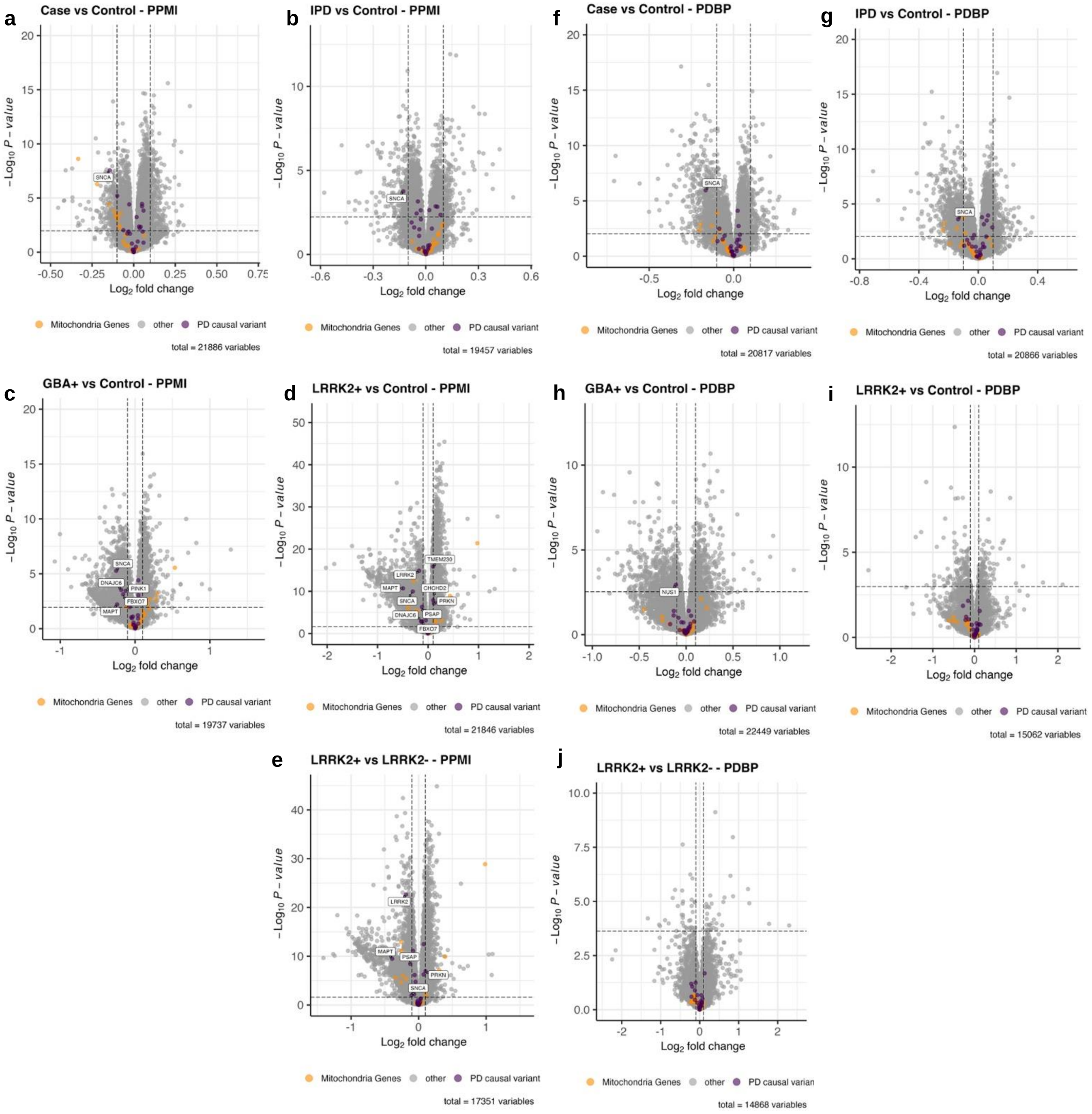


**Supplemental Figure 4 | Differential expression analysis with predicted neutrophil percentage correction split by study.** DE analyses were conducted with only PPMI samples. PD causal variants are colored purple, and mitochondrial genes are colored orange. Differentially expressed PD causal variants are additionally labeled by gene name. **a**, All PD cases were compared to all control samples based on the diagnosis at baseline. **b**, Idiopathic case samples with no SNCA+/LRRK2+/GBA+ mutation were compared to control samples with no PD-related mutations. **c,d,e,** Control samples with no PD-related mutations were compared to case samples with GBA+ (**c**) and LRRK2+ (**d**) mutations. LRRK2+ case vs LRRK2- case samples in PPMI were also compared (**e**). **f,g,h,i,j,** The same analyses were conducted, but this time with only PDBP sample. SNCA+ samples were only present in PPMI and were included in Figure 4.

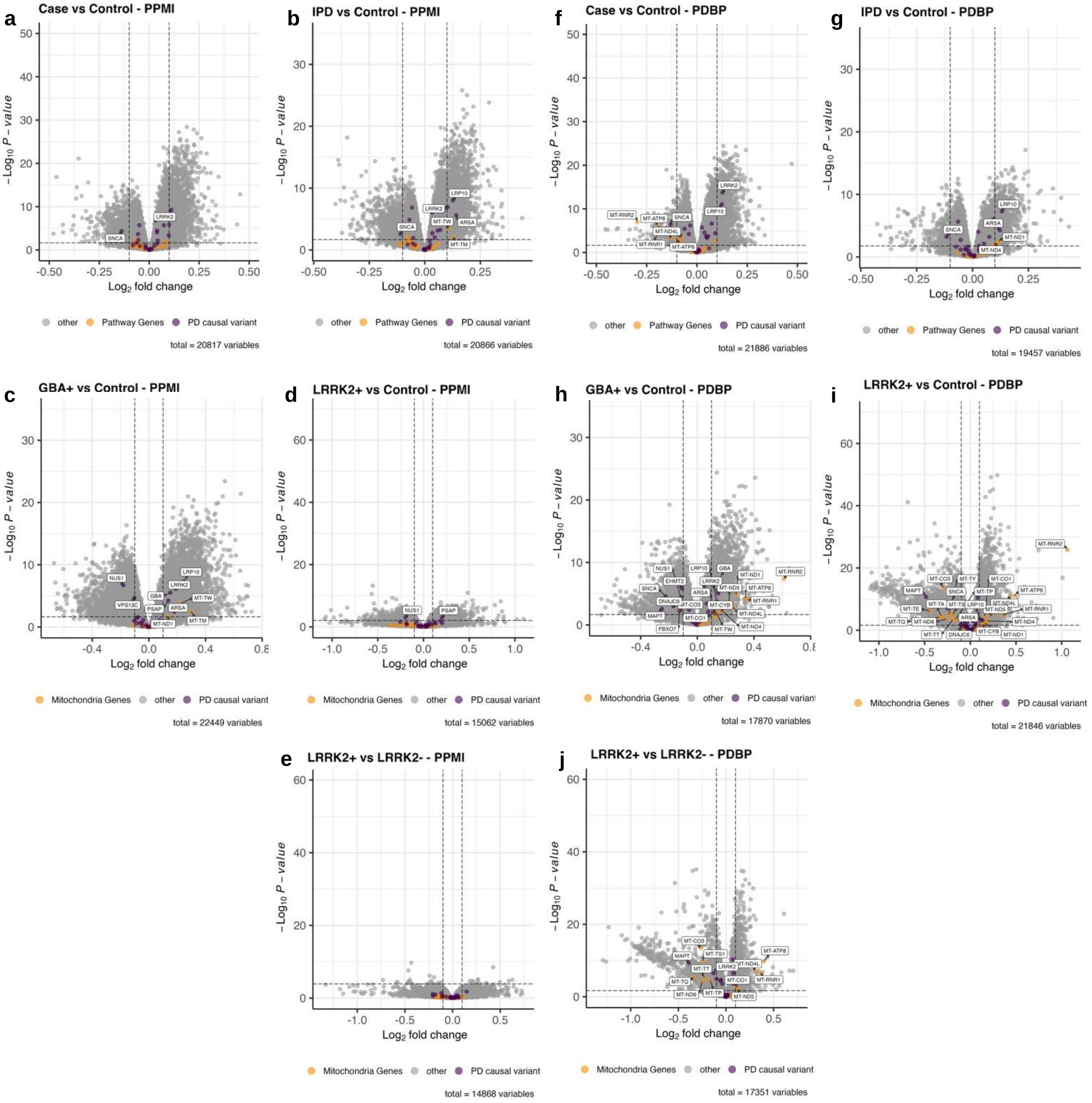


**Supplemental Figure 5 | Differential expression analysis without predicted neutrophil percentage correction split by study.** DE analyses were conducted with only PPMI samples but without including predicted neutrophil percentage in the covariate design. PD causal variants are colored purple, and mitochondrial genes are colored orange. Differentially expressed PD causal variants are additionally labeled by gene name. **a,** All PD cases were compared to all control samples based on the diagnosis at baseline. **b,** Idiopathic case samples with no SNCA+/LRRK2+/GBA+ mutation were compared to control samples with no PD-related mutations. **c,d,e,** Control samples with no PD-related mutations were compared to case samples with GBA+ (**c**) and LRRK2+ (**d**) mutations. LRRK2+ case vs LRRK2- case samples in PPMI were also compared (**e**). **f,g,h,i,j,** The same analyses were conducted, but this time with only PDBP sample. SNCA+ samples were only present in PPMI and were included in Supplemental Figure 2.

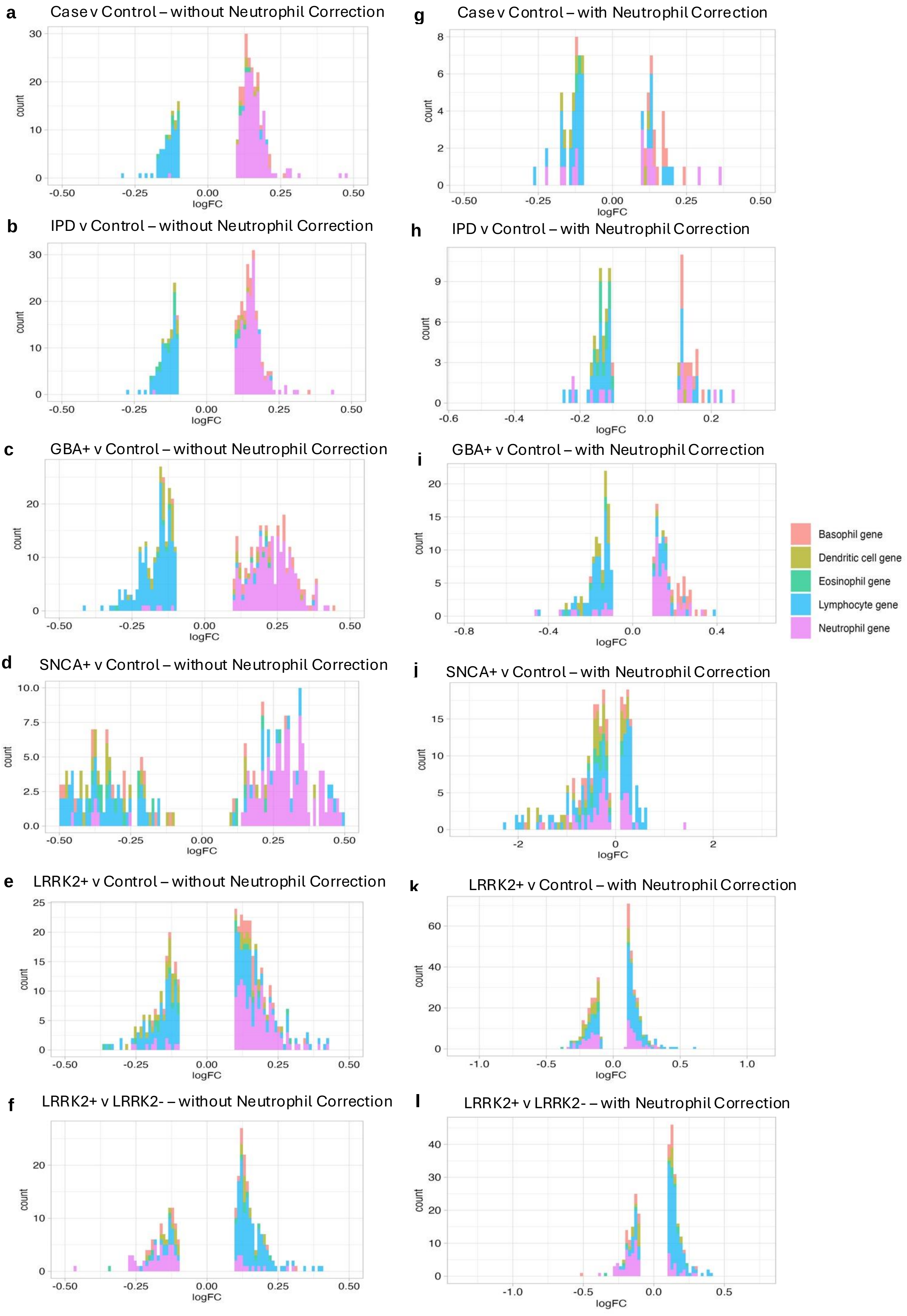


**Supplemental Figure 6 | Bar plots of differential expressed blood cell-enriched genes by PD cohort vs control comparison. a,b,c,d,e,f,** Blood cell-enriched genes that are differentially expressed in DE analyses without including predicted neutrophil percentage in covariate design. **g,h,i,j,k,l,** Blood cell-enriched genes that are differentially expressed in DE analyses with predicted neutrophil percentage in covariate design.

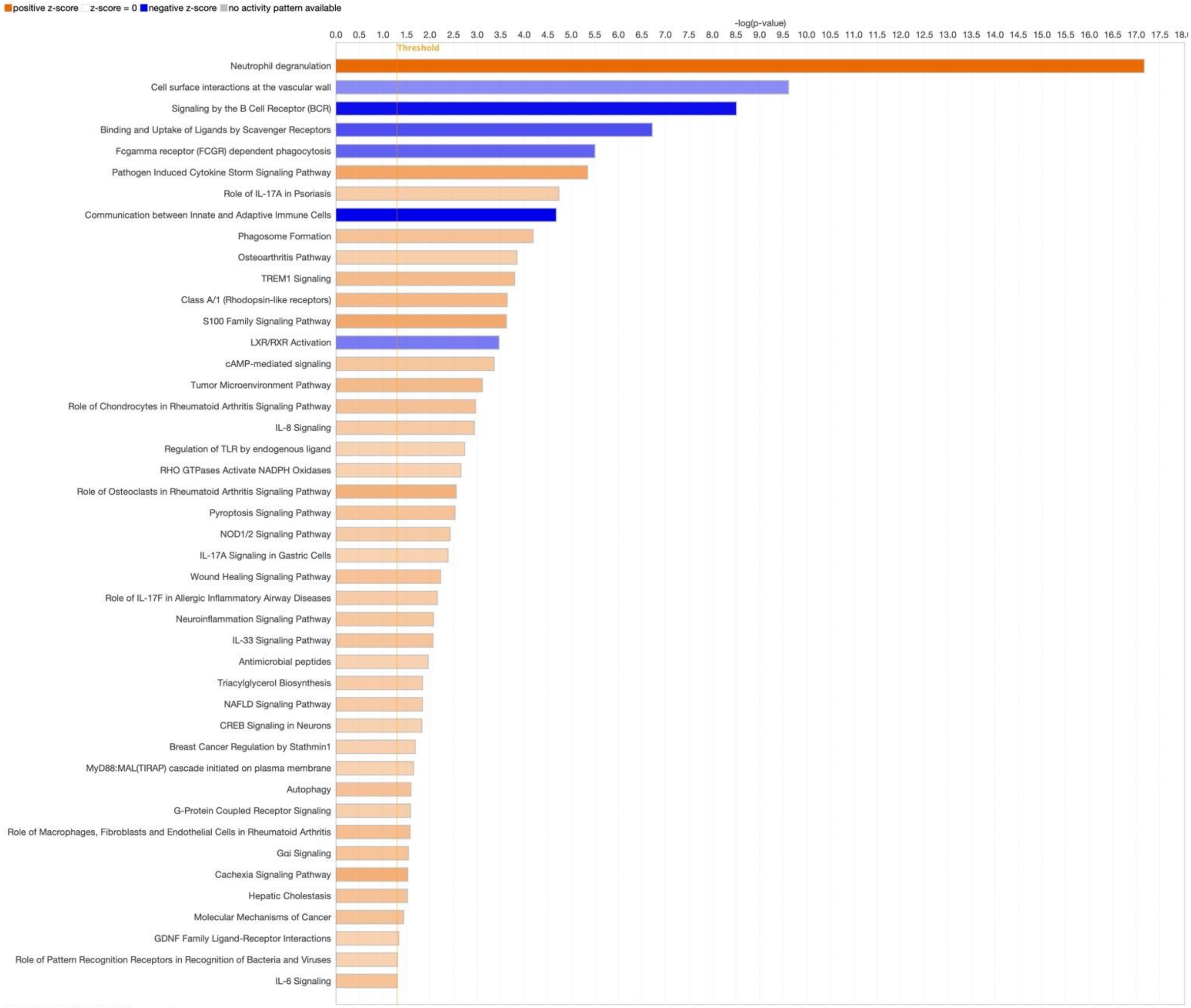


© 2000-2024 QIAGEN. All rights reserved.

**Supplemental Figure 7 | Differentially expressed pathways in PD case vs control in all samples without predicted neutrophil percentage correction.**

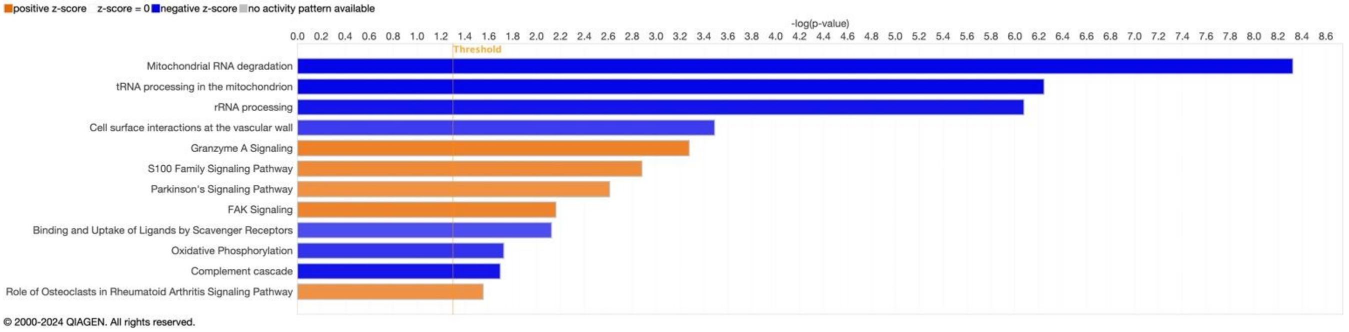


© 2000-2024 QIAGEN. All rights reserved.

**Supplemental Figure 8 | Differentially expressed pathways in PD case vs control in all samples with predicted neutrophil percentage correction.**

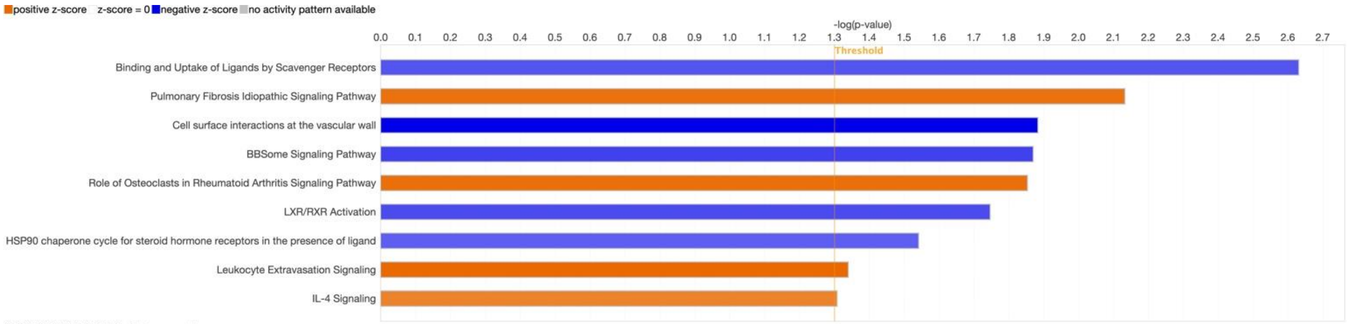


© 2000-2024 QIAGEN. All rights reserved.

**Supplemental Figure 9 | Differentially expressed pathways in IPD vs control in all samples with predicted neutrophil percentage correction.**

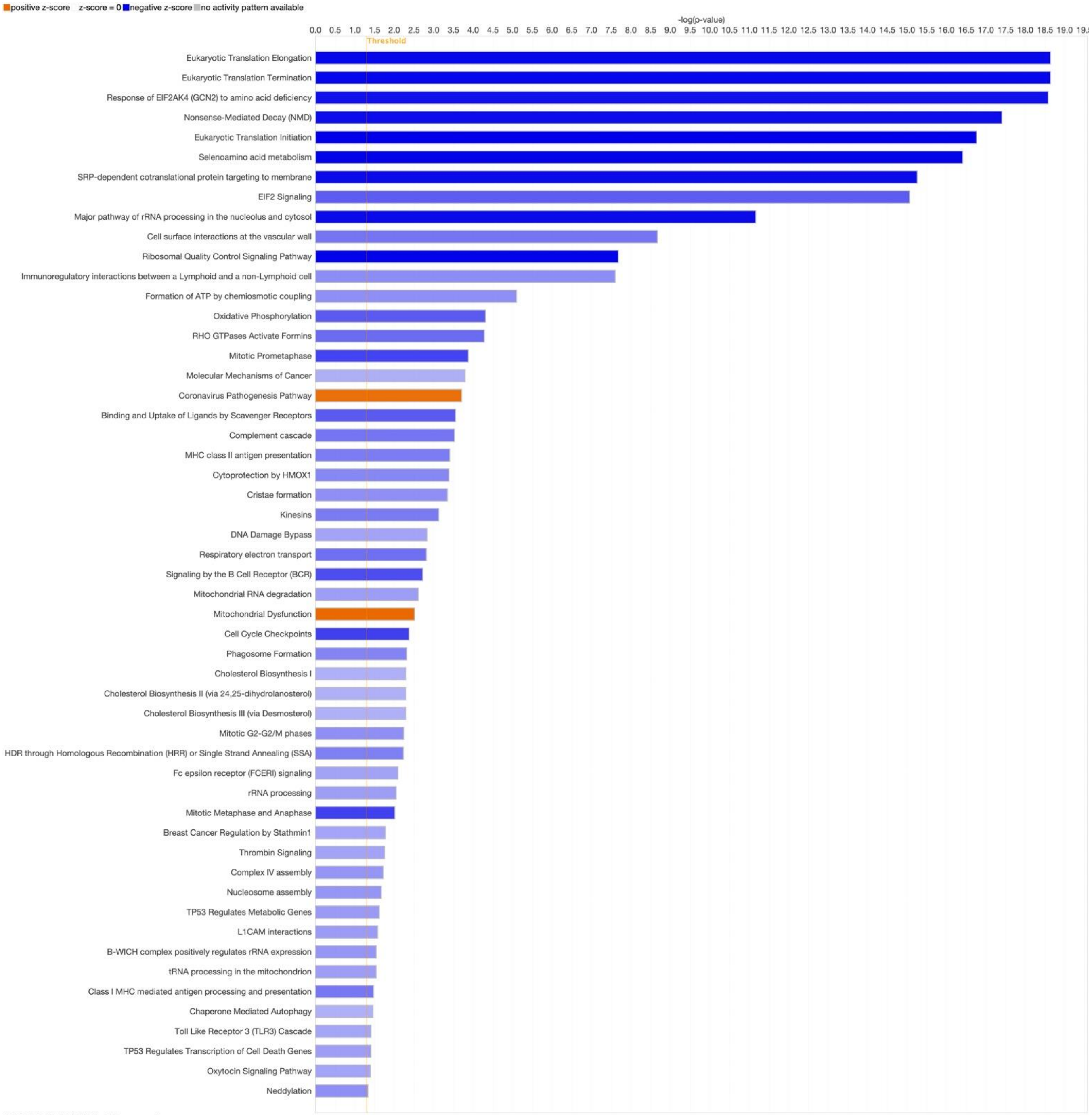


© 2000-2024 QIAGEN. All rights reserved.

**Supplemental Figure 10 | Differentially expressed pathways in GBA+ case vs control in all samples with predicted neutrophil percentage correction.**

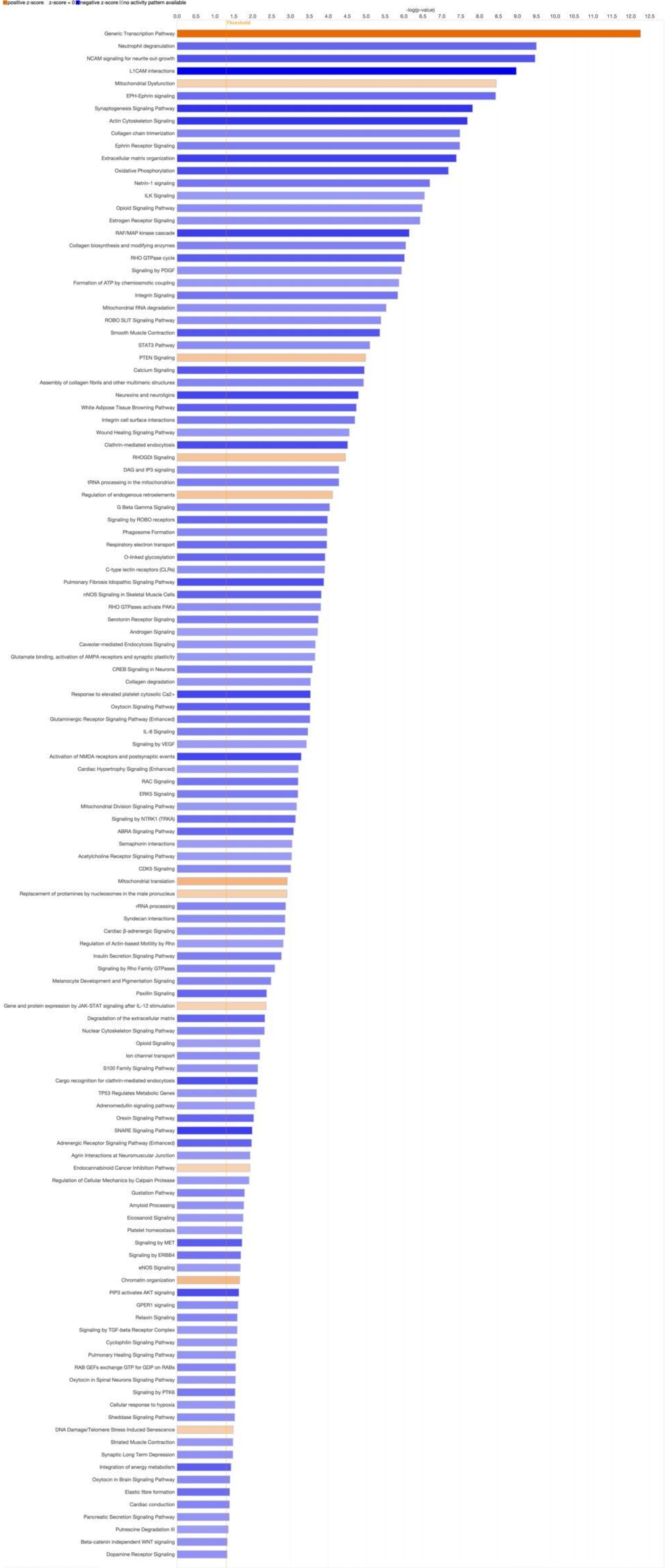


© 2000-2024 QIAGEN. All rights reserved.

**Supplemental Figure 11 | Differentially expressed pathways in SNCA+ case vs control in all samples with predicted neutrophil percentage correction.**

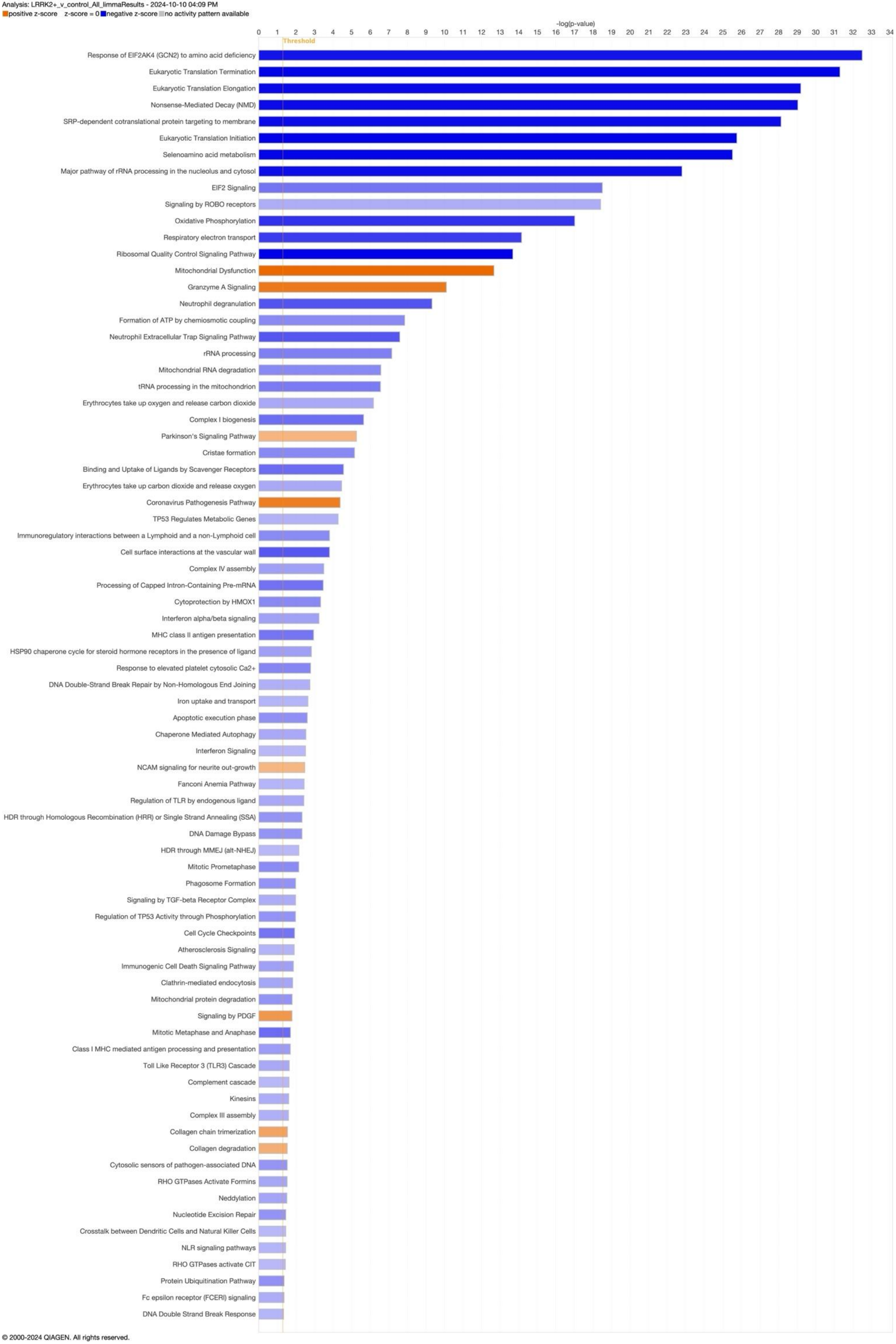


© 2000-2024 QIAGEN. All rights reserved.

**Supplemental Figure 12 | Differentially expressed pathways in LRRK2+ case vs control in all samples with predicted neutrophil percentage correction.**

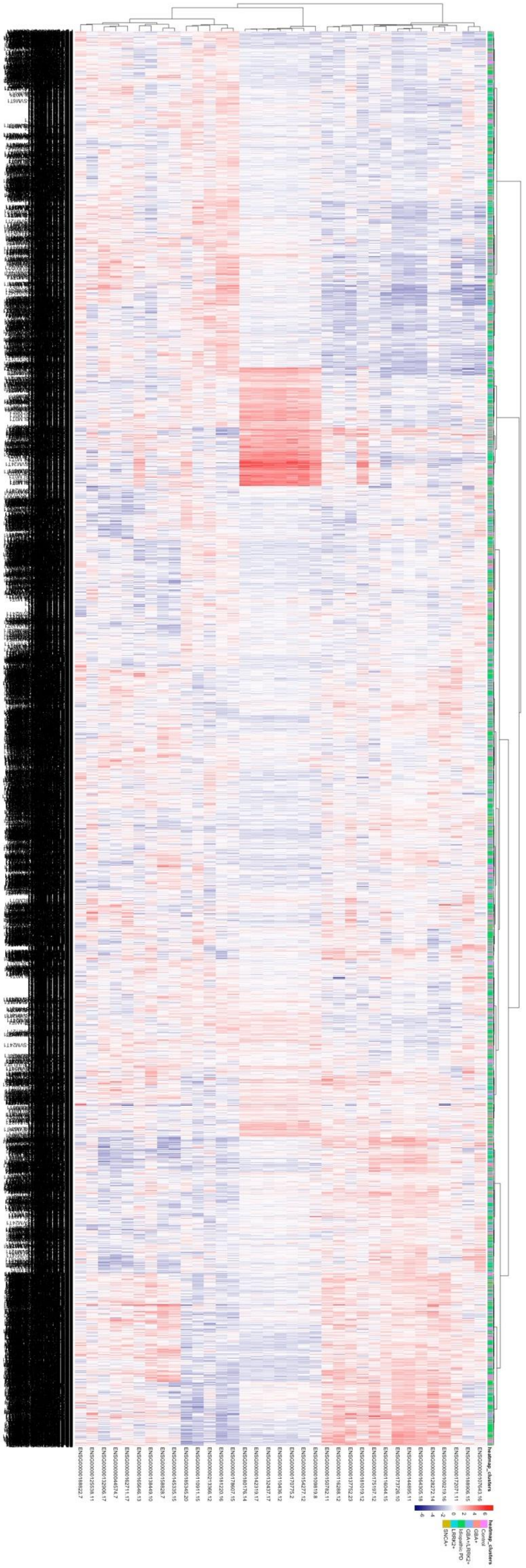

**Supplemental Figure 13 | Hierarchical clustering of PPMI and PDBP samples by 'Parkinson's Signaling Pathway' genes.**

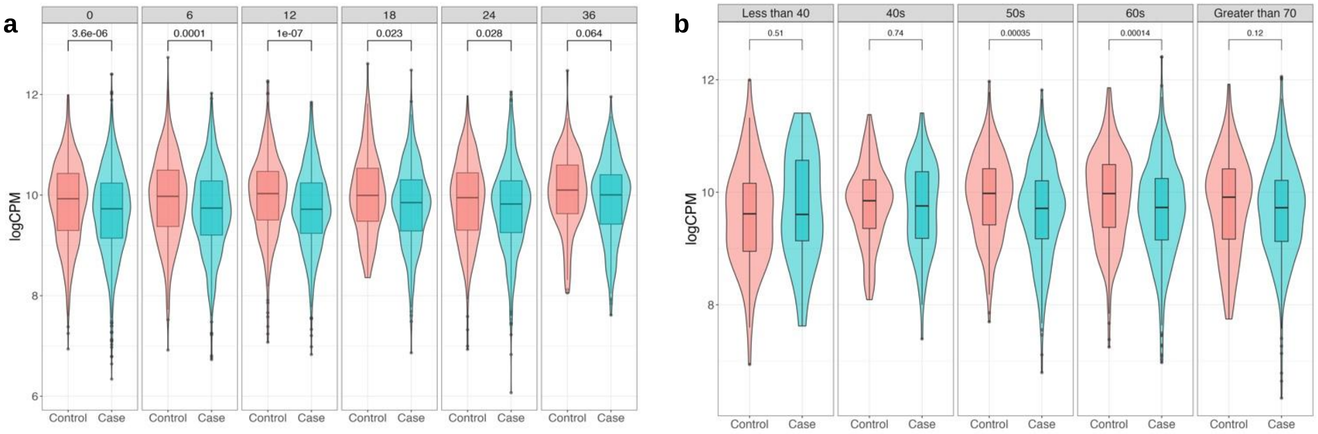


**Supplemental Figure 14 | SNCA expression plots by participant visit month and age at baseline sample.** SNCA gene counts were log(CPM) normalized and corrected for predicted neutrophil percentage. **a,** Violin plots of SNCA expression stratified by visit month. **b,** Violin plots of SNCA expression stratified by age at enrollment into PPMI or PDBP. A two-tailed Wilcoxon test was applied to compare case and control expression.

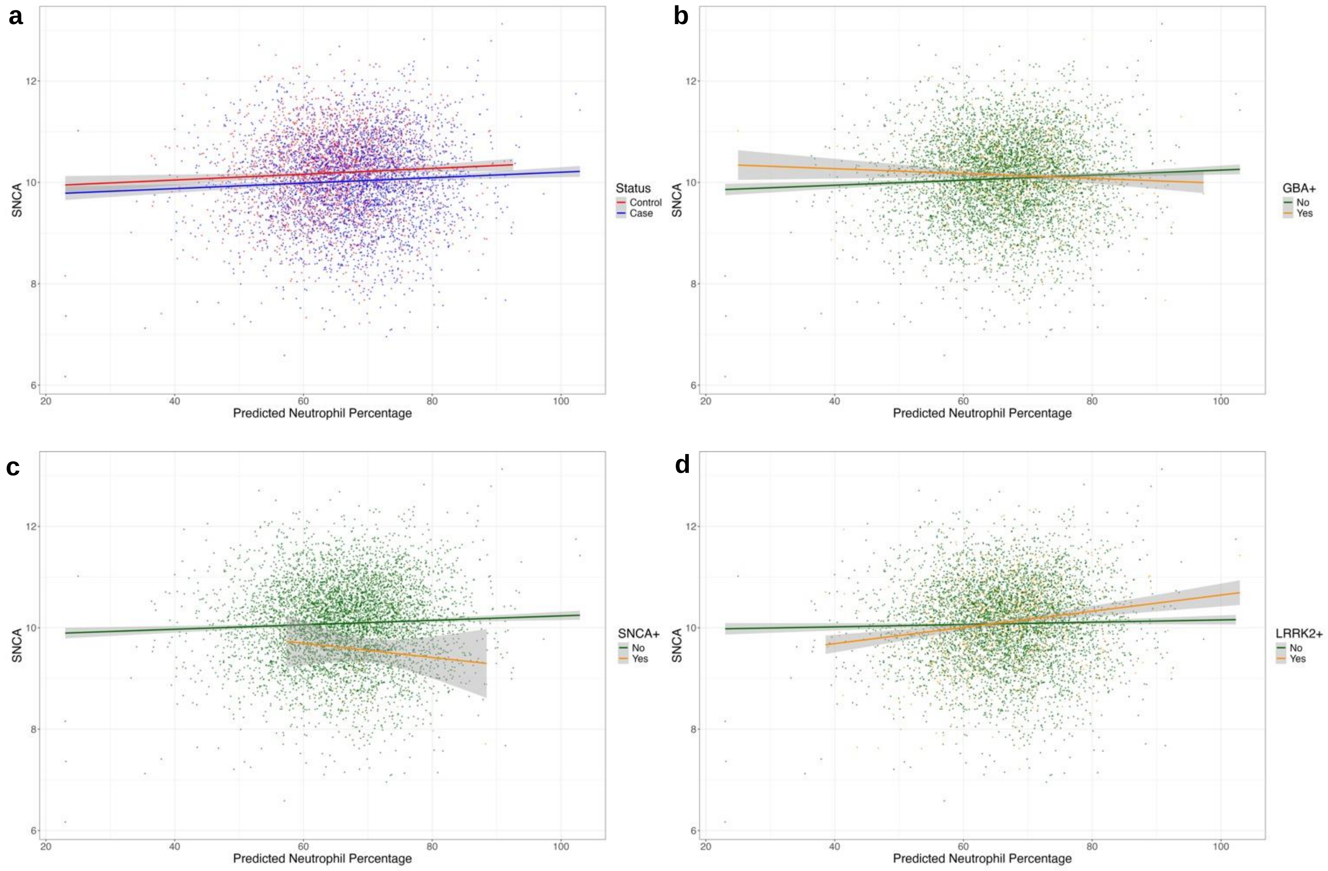


**Supplemental Figure 15 | SNCA expression by predicted neutrophil percentage.** SNCA counts were log(CPM) normalized for plotting. **a**, SNCA expression by predicted neutrophil percentage, colored by disease status (PD case and control). **b**, SNCA expression by predicted neutrophil percentage in GBA+ and GBA- samples. **c**, SNCA expression by predicted neutrophil percentage in SNCA+ and SNCA- samples. **d**, SNCA expression by predicted neutrophil percentage in LRRK2+ and LRRK2- samples.
